# Non-retroviral Endogenous Viral Element Acts as a Stable Antiviral Regulator Across Mosquito Life Stages and Generations

**DOI:** 10.64898/2026.07.22.739966

**Authors:** Yasutsugu Suzuki, Artem Baidaliuk, Yu Sekii, Irish Coleen A Asin, Pascal Miesen, Matthieu Prot, Louis Lambrechts, Etienne Simon-Loriere, Kozo Watanabe

## Abstract

Endogenous viral elements (EVEs) from non-retroviral RNA viruses are widespread in host genomes, yet their functional significance *in vivo* remains poorly understood. In mosquitoes, EVEs have been shown to suppress cognate viral replication through the PIWI-interacting RNA (piRNA) pathway. However, how this antiviral activity shapes virus-host dynamics beyond reducing viral replication remains unclear. Here, we established *Aedes aegypti* mosquito lines naturally infected with cell-fusing agent virus (CFAV) that differ in the presence or absence of a CFAV-derived EVE (CFAV-EVE1). Using this natural virus-host system, we showed that CFAV-EVE1 suppressed viral replication across sexes, tissues, developmental stages, and generations. This antiviral effect is initiated by EVE-derived primary piRNAs, as demonstrated in ovaries and heads. Despite reducing viral load, CFAV-EVE1 conferred no measurable benefits on mosquito survival or reproduction, nor did it eliminate CFAV from the naturally infected population under standard laboratory conditions. However, vertical transmission rates were reduced in EVE-containing mosquitoes and prolonged egg storage further reduced viral prevalence, suggesting that environmental stress may reveal EVE-mediated viral suppression. Together, these findings demonstrate that EVEs function as stable, heritable regulators of persistent virus-host interactions and highlight that ecological context may be critical for fully appreciating their functional significance.

## Introduction

Endogenous viral elements (EVEs) are viral sequences that have integrated into host genomes and are increasingly recognized as important contributors to physiological functions ^1^. While retroviral EVEs are well established in many mammalian lineages because retrovirus actively integrate their viral DNA into host genome as part of their life cycle, recent genomic studies have revealed that EVEs derived from non-retroviral RNA viruses are also widespread in diverse hosts ^2–7^. Despite their prevalence, we are only beginning to appreciate the biological significance of non-retroviral EVEs (nrEVEs), particularly in relation to virus infection. In case of retroviral EVEs, it has been demonstrated that these elements can inhibit viral replication through multiple mechanisms, including the expression of EVE-derived proteins or regulatory RNAs ^8–12^. In contrast, the antiviral functions of nrEVEs are still largely unexplored. Although large numbers of nrEVEs have been identified through bioinformatic analyses, especially in invertebrate genomes, experimental evidence directly linking nrEVEs to antiviral immunity remains limited. Addressing this gap requires a model system in which naturally circulating viruses and experimental manipulation can be combined.

Mosquitoes are one of the most widely used model organisms for studying nrEVEs. Several bioinformatic studies have revealed that the genomes of two major *Aedes* mosquito vectors, *Ae. aegypti* and *Ae. albopictus*, harbor hundreds of nrEVEs ^6,7,13,14^. These nrEVEs are frequently embedded within PIWI-interacting RNA (piRNA)-producing loci, a phenomenon that has been observed across diverse animal taxa ^6,15^. The piRNA pathway primarily play a role in suppressing expression of transposable elements, which can be deleterious to genome integrity in germlines ^16^. Notably, mosquitoes possess a piRNA pathway that is active in both germline and somatic tissues ^17,18^, making them a tractable model for investigating the functional interplay between EVEs, the piRNA pathway, and viral infections.

Taxonomically, most mosquito EVEs exhibit closest sequence similarity with mosquito-specific viruses (MSVs) ^7^, whereas EVEs closely related to arthropod-borne viruses (arboviruses) have not been reported. This bias is likely explained by the transmission ecology of MSVs, which are thought to be maintained in mosquito populations through vertical transmission ^19–22^, leading to frequent infection of germline cells and a higher probability of viral integration into the germline genome.

Importantly, the majority of mosquito nrEVEs are highly divergent from currently circulating MSVs, suggesting that they are derived from likely extinct viruses, thus precluding direct experimental testing of whether EVE-derived piRNAs can target cognate viral RNA and suppress replication. A notable exception to this is the MSV, cell-fusing agent virus (CFAV), for which we have previously identified a nrEVE in natural *Ae. aegypti* populations ^17^. Using this system, we demonstrated that a CFAV-derived EVE (named CFAV-EVE1) restricts CFAV replication in *Ae. aegypti*, particularly in the ovaries, through the piRNA pathway. However, whether and how this EVE-mediated antiviral activity influences mosquito and viral fitness remain unknown. One possibility is that suppressing viral replication in ovaries helps maintain normal oogenesis and egg quality, because high viral loads in ovarian tissues may impose physiological stress or interfere with proper oocyte development. Another non-mutually exclusive possibility is that reduced viral replication lowers the efficiency of vertical transmission, as decreased parental viral loads could limit the amount of virus passed to the progeny.

Experimental MSV infection in mosquitoes is often established by intrathoracic injection, because no reliable method exists to recapitulate natural MSV infection while maintaining control of the host genetic background. However, artificial infection routes frequently exhibits markedly reduced vertical transmission, likely because ovarian follicles are not infected under these conditions ^23,24^. Such aberrant tissue tropism may therefore fail to recapitulate natural infection phenotypes. In addition, intrathoracic injection causes physical damage, and viral replication kinetics may differ from those observed during natural infection. Consequently, accurate evaluation of EVE-mediated antiviral functions in MSV vertical transmission and mosquito reproductive traits requires the use of mosquitoes naturally infected with MSVs.

Here, we establish genetically matched *Ae. aegypti* mosquito lines that are naturally infected with CFAV and differ in the presence or absence of CFAV-EVE1. This allows us to test whether EVE-mediated antiviral activity functions as a heritable trait that influences mosquito reproductive fitness and viral vertical transmission under natural infection conditions. By examining EVE function across sexes, the mosquito life cycle, and over multiple generations, our work provides a broad evaluation of the role of nrEVEs in regulating persistent viral infections *in vivo*.

## Results

### Generation of naturally CFAV-infected *Aedes aegypti* mosquito lines with or without CFAV-EVE1

To investigate the antiviral role of CFAV-EVEs in a natural infection context, we established *Ae. aegypti* mosquito lines naturally infected with CFAV, either carrying or lacking CFAV-EVE1, by repeated crosses between different mosquito strains (Figure 1A, Figure S1C). For this purpose, we made use of a naturally CFAV-infected *Ae. aegypti* colony from Vietnam, which we confirmed to encode two previously described CFAV-EVEs ^17,25^, namely CFAV-EVE1 and CFAV-EVE2, respectively (Figure S1A). CFAV-infected females from this strain were first crossed with CFAV-free males from a CFAV-EVE1 knockout (KO) line previously generated using CRISPR/Cas9 genome editing ^17^ (Figure S1C). The resulting female progeny were then backcrossed with CFAV-EVE1 KO males for three successive generations, maintaining the direction of the cross (Vietnam colony-derived females × KO males) throughout. After the third backcross (fourth cross in total), individuals that retained natural CFAV infection and lacked CFAV-EVE1 and CFAV-EVE2, as confirmed by PCR, were selected. These females were then crossed with CFAV-free males homozygous for CFAV-EVE1 (+/+) in the absence of CFAV-EVE2, which represent a sister line of the KO line, also generated in our previous study ^17^. From the resulting heterozygous (+/−) progeny, we generated homozygous CFAV-EVE1 (+/+) and (−/−) individuals through sibling crosses (Figure 1A). F2 individuals were genotyped for CFAV-EVE1 and separated into cages according to genotype. Using F3 progeny, we established three biological replicate lines per genotype. In the subsequent F4 generation, we confirmed the expected CFAV-EVE1 genotypes (n=32 in each). We also verified the genetic diversity of CFAV in the parental mosquitoes at the stage of the 4^th^ cross (Figure S2). We recovered nearly full CFAV genomes from 12 out of 14 individual larvae. Intrahost single nucleotide variant frequencies were well below 25% in most samples except for two variants with higher frequencies (Figure S2A). Among the 12 consensus sequences, we observed only 8 mutations, from which 4 mutations in 4 different sequences were non-synonymous (Figure S2B). Importantly, none of the consensus mutations or the two high-frequency intrahost variants fell in the regions corresponding to CFAV-EVE1. Overall, the diversity in the colony is relatively low, given that the closest sequence to our sequences from Vietnam is from Cambodia (LR694077) and it is ∼2% different from our sequences (∼200 mutations).

**Figure 1.**
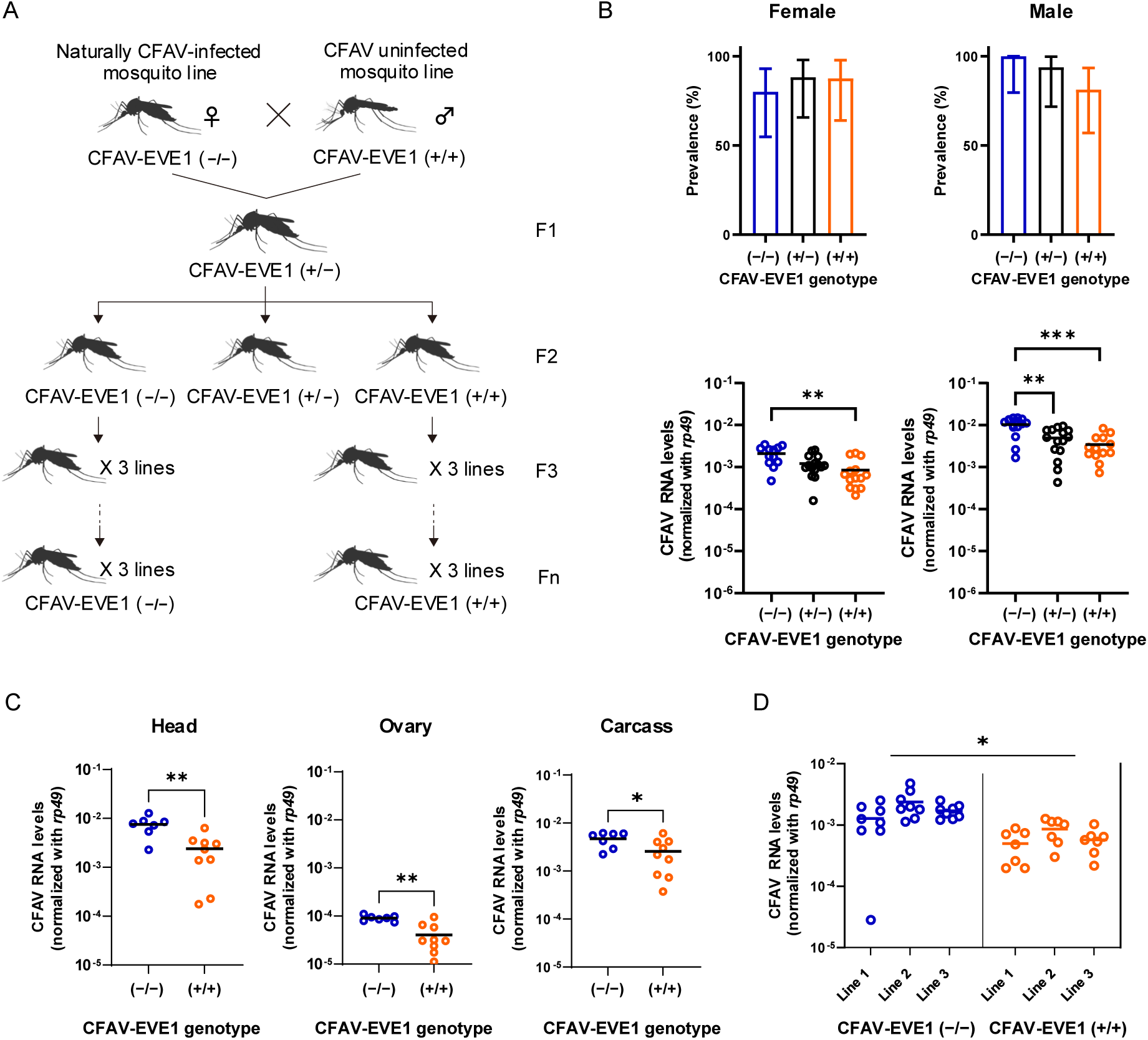
Effect of the CFAV-EVE1 genotype on CFAV infection in mosquitoes. (A) Schematic illustration of the experimental design used to establish *Ae. aegypti* mosquito lines naturally infected with CFAV and devoid of (−/−) or homozygous (+/+) for the CFAV-EVE1. CFAV-EVE1 (+/+) female naturally infected with CFAV were crossed with uninfected CFAV-EVE1 (−/−) males to generate mosquito lines naturally infected with CFAV and either CFAV-EVE1 (−/−) or CFAV-EVE1 (+/+). Each mosquito line was maintained for several generations (F0–Fn) in three independent cages. (B) CFAV prevalence and viral RNA levels in adult mosquitoes. The percentage of CFAV-positive individuals was measured separately in females and males for each genotype. Viral RNA levels were quantified by RT-qPCR and normalized with the housekeeping gene *rp49*. Only CFAV-positive individuals are included in the RNA level analysis. (C) CFAV RNA levels in head, ovary, and carcass were quantified by RT-qPCR and normalized to *rp49* transcripts. (D) Replicate line-specific comparison of CFAV RNA levels between CFAV-EVE1 genotypes. Viral RNA levels were quantified in mosquitoes from three independent lines. Data represent individual mosquitoes (B and D) or pools of three individuals (C). Statistical significance represented by asterisks was assessed with Fisher’s exact (B, prevalence), Wilcoxon rank-sum (B, C, RNA levels) pairwise tests and nested t-test (D) (*P <0.05; **P ≤0.01; ***P ≤0.001).

### CFAV replication is suppressed by CFAV-EVE1 in naturally infected mosquitoes

To assess the impact of CFAV-EVE1 on CFAV replication in the naturally infected mosquitoes, we measured CFAV RNA levels in female and male adults with different genotypes: CFAV-EVE1 (−/−), (+/−) and (+/+) (Figure 1B). The proportion of CFAV-positive individuals ranged from 80% to 100% across lines and sexes and pairwise differences between genotypes were insignificant. Interestingly, only in males, an increase in the number of CFAV-EVE1 positive alleles modestly reduced the infection proportion (GLM *P =* 0.04404, Figure 1B, Table S1). In both sexes, although not all genotype comparisons reached statistical significance, CFAV RNA levels in CFAV-positive individuals showed a decreasing trend from CFAV-EVE1 (−/−) to (+/−) to (+/+) mosquitoes (Figure 1B, Spearman’s ρ = -0.623, *P* < 0.0001 and ρ = -0.558, *P =* 0.000152 in males and females, respectively, Table S2). We further quantified CFAV RNA levels in the ovaries, heads, and carcasses of naturally infected female mosquitoes from CFAV-EVE1 (−/−) and (+/+) lines. CFAV RNA levels were significantly lower in CFAV-EVE1 (+/+) mosquitoes compared to CFAV-EVE1 (−/−) mosquitoes across tissues (*P* = 0.03335, Table S3 even though the overall levels varied by tissue as well (*P* < 0.0001, Table S3) (Figure 1C). No significant interaction effect was observed between CFAV-EVE1 genotype and tissue (*P* = 0.48337, Table S3) (Figure 1C). These results suggest that CFAV-EVE1 contributes to antiviral activity across multiple tissues in naturally infected mosquitoes. Consistent patterns in viral suppression were observed across the three independent lines for each genotype (Figure 1D).

### CFAV-EVE1 is required for ping-pong amplification of viral piRNAs in naturally infected mosquitoes

Our previous study demonstrated that the CFAV-EVE1-mediated antiviral mechanism operates via the piRNA pathway. Therefore, we assessed whether CFAV-EVE1 absence results in alterations of the cognate viral piRNA populations in the naturally infected mosquitoes. We sequenced the small RNAs of CFAV-infected CFAV-EVE1 (+/+) and (−/−) mosquitoes, focusing on ovary and head tissue to independently analyze germline and somatic tissues, respectively. Overall, we generated twelve small RNA libraries representing three independent biological replicates for both tissues in both genotypes. These libraries had the expected size distribution of small RNAs with a prominent peak at 22 nucleotides (nt) representing cellular microRNAs (Figure S3). Moreover, the libraries prepared from ovary samples showed an abundant population of 25-30 nt reads, indicative of efficient piRNA production in the mosquito germline.

We next focused on CFAV-derived piRNAs comparing CFAV-EVE1 (+/+) and (−/−) mosquitoes. Viral piRNA production was observed both in ovary and head tissue. In EVE1 (+/+) mosquitoes, a discrete population of piRNA-sized small RNAs was produced from both genomic strands that mapped to the viral genome near the NS2 gene in both ovaries (Figure 2A, B) and heads (Figure S4A, B). These piRNAs map to a 686-nt genomic segment that is ∼96% identical to CFAV-EVE1. Strikingly, in CFAV-EVE1 (−/−) mosquitoes, this piRNA population at the cognate position of the CFAV-EVE1 is strongly diminished. Antisense piRNAs are essentially absent, and sense piRNAs are strongly reduced and do not map to the discrete NS2 region of the CFAV genome in both ovaries (Figure 2C, D) and heads (Figure S4C, D). Instead, piRNA populations in CFAV-EVE1 (−/−) mosquitoes are dominated by another highly discrete piRNA species that maps to the 3’ end the viral genome. The exact nature of this piRNA-sized small RNA species and its biogenesis are unknown. The disproportionate production of piRNAs mapping to the CFAV-EVE1 sequence in CFAV-EVE1 (+/+) compared to CFAV-EVE1 (−/−) mosquitoes is in line with our previously proposed model of EVE-triggered piRNA amplification via the ping-pong loop ^17^. This is also supported by a strong enrichment of 10-nt overlap in overlapping piRNAs derived from complementary RNA strands in both ovaries (Figure 2E) and heads (Figure S4E). The typical enrichment of an adenine at position 10 of sense piRNAs is visible but does not stand out compared to other, primarily U nucleotide biases along the piRNA sequences in both ovaries (Figure 2G) and heads (Figure S4G). This might be due to the limited sequence space of the EVE region which may increase noise in the analysis of sequence motifs. Most strikingly, the 10-nt overlap signature is absent in CFAV-EVE1 (−/−) mosquitoes, indicating that ping-pong amplification of viral piRNAs does not occur in the absence of the EVE (Figure 2F and Figure S4F). Overall, these data confirm the model in which EVE-derived primary piRNAs trigger ping-pong amplification of the cognate viral region, a process that is effectively ablated in mosquitoes lacking the integrated viral sequence.

**Figure 2.**
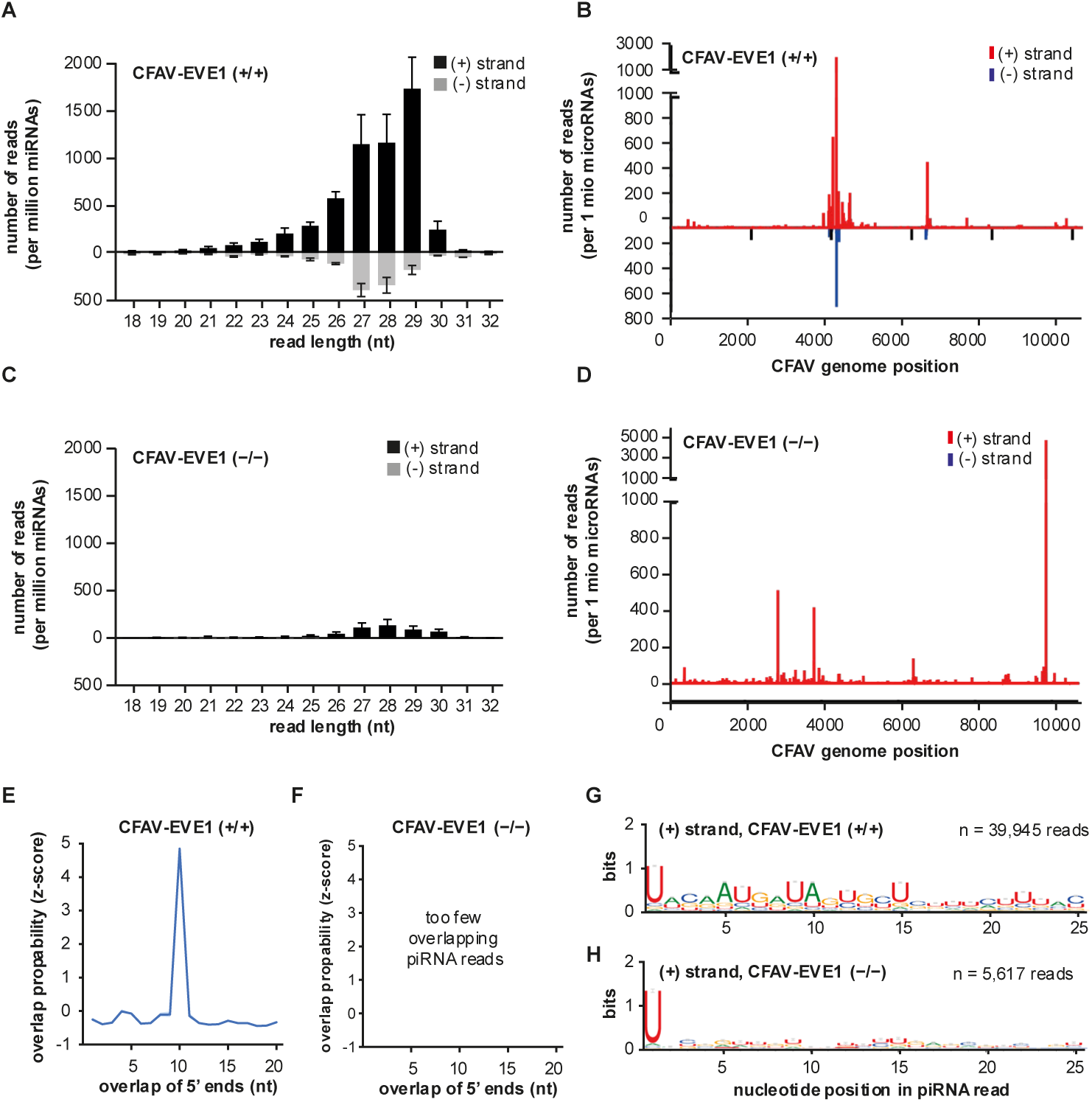
CFAV-EVE1-derived piRNAs interact with CFAV RNA in naturally infected mosquito ovaries. Size distribution (A, C), genome-wide distribution of piRNA-like small RNAs (25–30 nt) (B, D), overlap probability analysis of 25–30 nt small RNAs (E, F), and sequence logo analysis depicting nucleotide bias of 25–30 nt small RNAs (G, H) mapping to the CFAV genome in ovaries of CFAV-EVE1 (+/+) and (−/−) mosquitoes, respectively.

### CFAV-EVE1 does not confer a reproductive or fitness advantage in naturally infected mosquitoes

To test our first hypothesis that CFAV-EVE1 improves the mosquitoes’ reproductive success by suppressing CFAV replication in ovaries, we compared the fecundity (number of eggs laid) and fertility (larval hatching rate) of individual females between CFAV-EVE1 (+/+) and (−/−) lines. The experiments were repeated three times. Overall, no notable differences were observed in either fecundity or fertility between the two genotypes (Figure S5).

Because reproductive output was unaffected, we next assessed survival, another major component of fitness. CFAV RNA levels were lower in whole bodies of both female and male CFAV-EVE1 (+/+) mosquitoes than in CFAV-EVE1 (−/−) mosquitoes. We therefore asked whether viral suppression in presence of the EVE was associated with increased adult survival in either sex. Survival analysis indicated no difference in male mosquito survival between the two mosquito lines (mixed effects Cox model, *P* = 0.641, Table S4) (Figure S6). CFAV-EVE1 (−/−) females had a marginally significant survival advantage (mixed effects Cox model, *P* = 0.0235, Table S5). This effect reflects a slight relative decline in survival of CFAV-EVE1 (+/+) females after day 25 post adult emergence (Figure S6). These results suggest that although CFAV replication is suppressed in naturally infected CFAV-EVE1 (+/+) mosquitoes, this does not substantially impact reproductive output or survival under standard laboratory conditions.

### CFAV-EVE1 limits vertical transmission of CFAV

Next, we examined whether CFAV-EVE1 inhibits the vertical transmission of CFAV, potentially leading to its elimination from the population. To address this, we monitored both CFAV prevalence and CFAV RNA levels by RT-qPCR across multiple generations (Figure 3). Interestingly, in the CFAV-EVE1 (−/−) line females, prevalence gradually increased from 87.5% at generation F1 to 100% at generation F4 and remained at that level in all subsequent generations examined (Figure 3A). In contrast, the CFAV-EVE1 (+/+) line showed a sharp drop in prevalence from generation F2 to F3, declining by approximately 20%, followed by a rebound to 88% at F4. Between generations F2 and F3 we observed the opposite changes in prevalence in the two CFAV-EVE1 homozygous genotypes. We did not observe any significant pairwise differences between the genotypes either in F2 (*P* = 0.3781) or F3 (*P* = 0.07226). Nor did the prevalence differences between F2 and F3 reach significance in CFAV-EVE1 (+/+) (*P* = 0.2347566) or CFAV-EVE1 (−/−) (*P* = 1). However, in a model that included both generations and both genotypes, we observed a significant generation × CFAV-EVE1 interaction effect (*P* = 0.02126, Table S6), possibly capturing the jump in prevalence in CFAV-EVE1 (−/−) from F2 to F3 and the drop in CFAV-EVE1 (+/+). Modeling the change to the following generation F4, the increase in prevalence from generation F3 to F4 was significant (*P* = 0.027576, Table S7), accounting for the genotype effect, which was also significant in these two generations (*P* = 0.004442, Table S7). In subsequent generations (F4–F10 for females and F5–F10 for males), the prevalence remained relatively stable (no significant generation effect), fluctuating between 67% and 88% (females) and 75% – 83% (males) in CFAV-EVE1 (+/+) (Figure 3A, B). While the difference in prevalence between CFAV-EVE1 (+/+) and (−/−) genotypes across generations was significant (CFAV-EVE1 effect *P* < 0.0001 in females, Table S8 and *P* < 0.0001 in males, Table S9), CFAV was not eliminated from the population. These results indicate that CFAV-EVE1 partially limits vertical transmission, leading to a transient reduction in CFAV prevalence without complete viral loss.

To examine whether the antiviral effect of CFAV-EVE1 is maintained over generations, we compared CFAV RNA levels between CFAV-EVE1 (+/+) and (−/−) lines across multiple generations in naturally infected individuals. In both sexes, CFAV RNA levels were consistently lower in the CFAV-EVE1 (+/+) line compared to the (−/−) line in most generations examined, confirming a robust antiviral effect of CFAV-EVE1 (Figure 3C, D; pairwise *P* < 0.05 at all generations except F8 in females). Concordantly with pairwise differences, the overall negative effect of CFAV-EVE1 (+/+) was observed in both females (F3–F10, *P* < 0.0001, Table S10) and males (F5–F10, *P* < 0.0001, Table S11). And the overall effect of the generation was significant (both *P* < 0.0001, Tables S9, S10) while CFAV-EVE1 by generation interaction effect was not significant. These results indicate that CFAV-EVE1 maintains a stable antiviral effect across generations in naturally infected mosquitoes.

**Figure 3.**
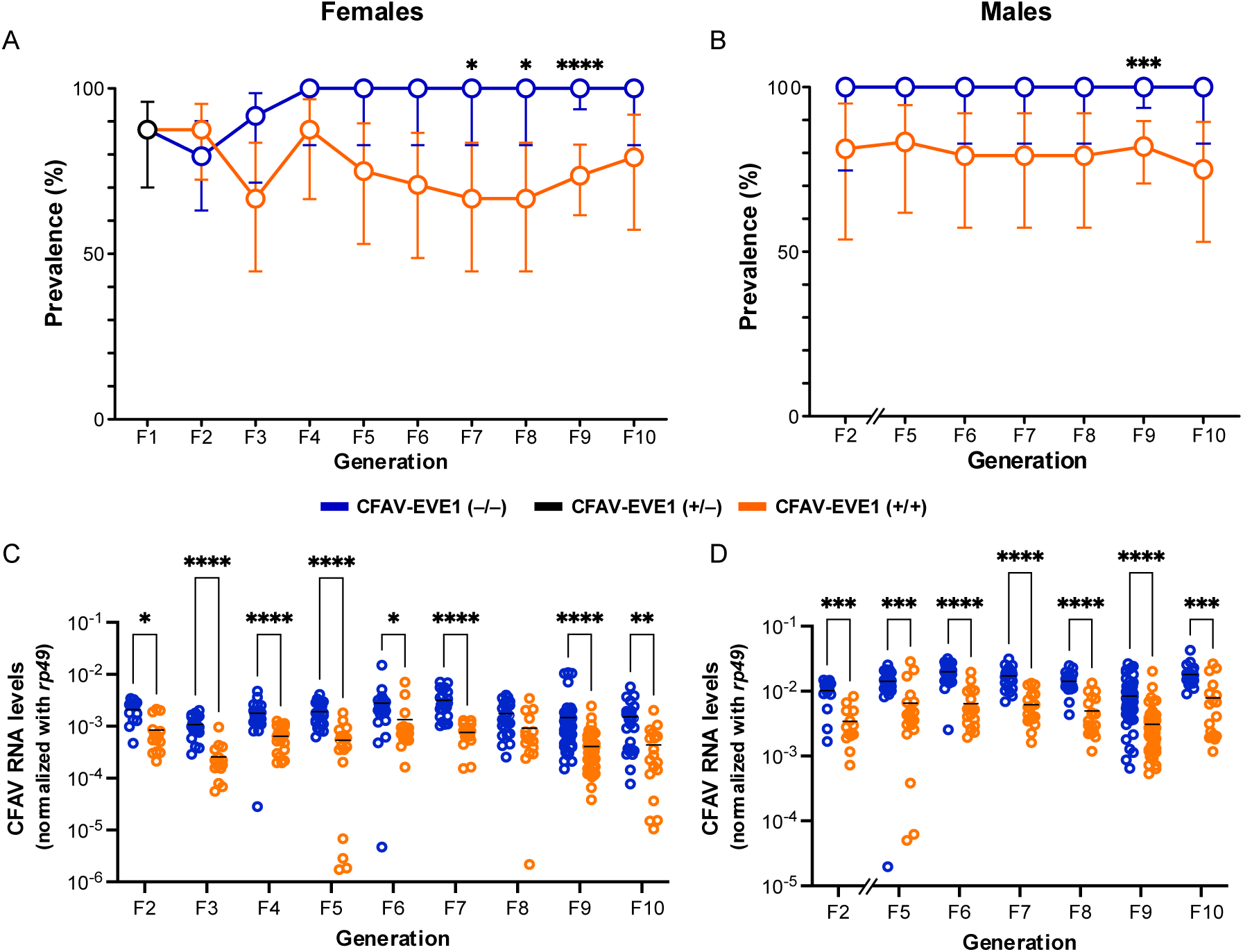
Effect of CFAV-EVE1 on CFAV infection across generations. CFAV infection prevalence across generations in CFAV-EVE1 (+/+) and (−/−) mosquitoes: (A) females and (B) males. Prevalence is based on data combined from all three replicate lines. Error bars represent 95% confidence intervals. CFAV RNA levels in CFAV-positive (C) adult females and (D) males across generations, normalized to *rp49* transcripts. Each dot represents an individual mosquito. Statistical significance represented by asterisks was assessed with Fisher’s exact (A, B), Wilcoxon rank-sum (C, D) pairwise tests (*P ≤0.05; **P ≤0.01; ***P ≤0.001; ****P ≤0.0001.

### CFAV-EVE1-mediated antiviral mechanism functions throughout mosquito life stages

To further explore the biological relevance of EVE-mediated antiviral effects, we examined whether CFAV replication is limited by CFAV-EVE1 during developmental stages prior to adulthood. All pooled samples of eggs and first-instar (L1) larvae from both CFAV-EVE1 genotypes tested positive for CFAV RNA. Starting from third-instar larvae (L3), CFAV prevalence was consistently lower in CFAV-EVE1 (+/+) compared to CFAV-EVE1 (−/−) individuals (Figure 4A). Although the pairwise reduction in CFAV prevalence only reached statistical significance at L3, the overall genotype effect across all life stages (coupled with sex in one variable) was statistically significant (*P* < 0.0001, Table S12), while the effect of the life stage and sex (as one variable) was not significant. In infected individuals, CFAV RNA levels were also significantly lower in the CFAV-EVE1 (+/+) line compared to the CFAV-EVE1 (−/−) line at most of developmental stages and two sexes (Figure 4B). Interestingly, no statistically significant pairwise difference was observed at the L3 stage, likely due to the reduced infection rate at this stage. We also observed that the negative effect of CFAV-EVE1 (+/+) genotype on CFAV RNA levels was much stronger in pupae and one-day-old (D1) females, which was also supported by a significant CFAV-EVE1 × life stage (merged with sex in one variable) interaction effect (*P* = 0.008363, Table S13; only pupae and adults included).

**Figure 4.**
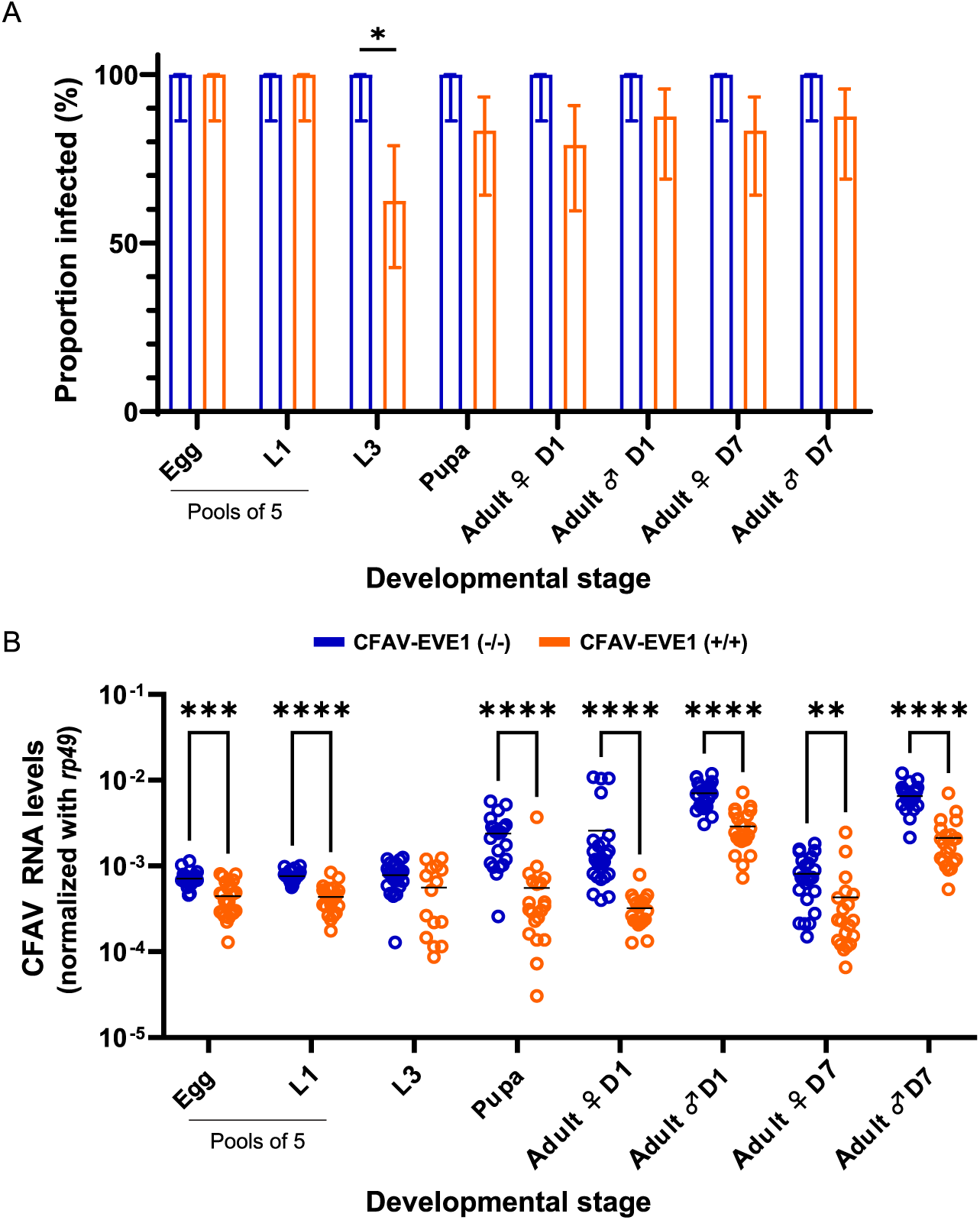
CFAV-EVE1 suppresses CFAV replication across mosquito developmental stages. (A) CFAV infection prevalence and (B) CFAV RNA levels across developmental stages (egg, 1st instar (L1), 3rd instar (L3), pupa, and adult) in mosquito lines carrying CFAV-EVE1 (+/+) or (−/−). In (B), each dot represents an individual mosquito, except for egg and L1 stages where each dot represents a pool of five individuals. CFAV RNA levels were normalized to rp49 transcripts. Statistical significance represented by asterisks was assessed with Fisher’s exact (A) Wilcoxon rank sum (B) pairwise tests (*P ≤0.05; **P ≤0.01; ***P ≤0.001; ****P ≤0.0001).

Unexpectedly, we observed that prolonged egg storage for 4 months prior to hatching, compared with the usual storage period of approximately 1 month, resulted in a marked reduction in CFAV prevalence among the resulting individuals (Figure 5A). The negative effect of the presence of CFAV-EVE1 on CFAV detection and/or RNA levels remained significant at most life stages and in both sexes (Figure 5A, B). To further evaluate the impact of the prolonged egg storage, we examined how CFAV prevalence and RNA levels varied with CFAV-EVE1 genotype and egg-storage conditions by analyzing adult female mosquitoes from two batches derived from stored eggs at generations F4 and F5 (Figure 5C, D and Figure S7A, B). Both CFAV prevalence and RNA levels were significantly lower in CFAV-EVE1 (+/+) mosquitoes (both *P* < 0.0001; Tables S14 and S15, respectively) or following extended egg storage (*P* = 0.000264 and *P* < 0.0001; Tables S14 and S15, respectively). The effects were reproducible in both generations (*i.e*., the effect of the generation was not significant). Although the interaction between CFAV-EVE1 genotype and egg storage was not statistically significant, the reduction in CFAV prevalence associated with the CFAV-EVE1 (+/+) genotype was more pronounced after prolonged egg storage. Together, these results indicate that CFAV-EVE1 exerts antiviral effects throughout mosquito development, which can be further enhanced under environmental stress such as prolonged egg storage.

**Figure 5.**
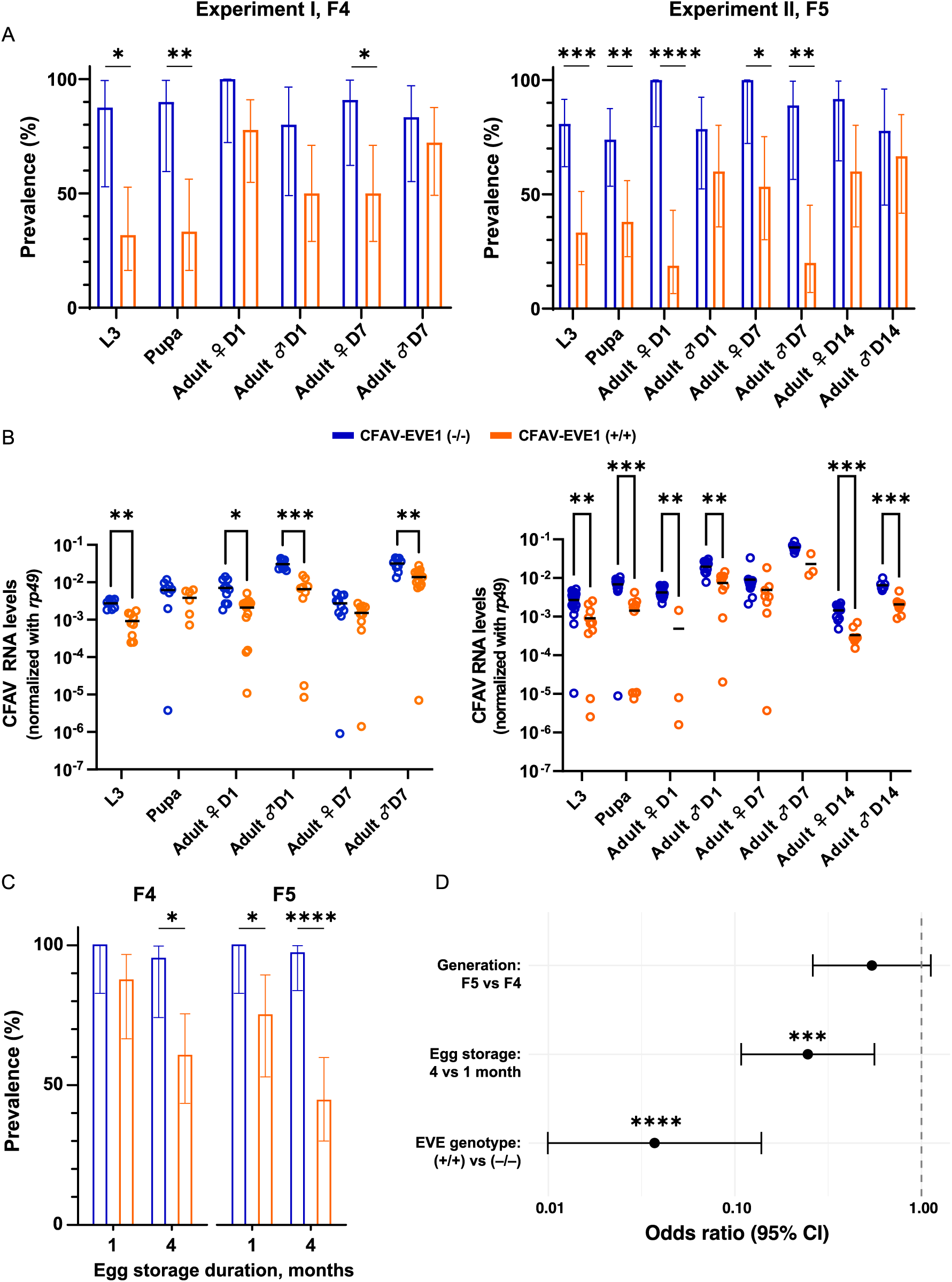
Effect of prolonged egg storage on CFAV dynamics between CFAV-EVE1 genotypes. (A) CFAV infection rate and (B) RNA levels across developmental stages (3rd instar (L3), pupa, and adult) in mosquitoes derived from eggs subjected to prolonged storage for 4 months, shown for two independent experiments in CFAV-EVE1 (+/+) and (−/−) lines. CFAV RNA levels were normalized to rp49 transcripts. Each dot in (B) represents an individual mosquito. (C) Side-by-side comparison of CFAV infection rate between CFAV-EVE1 genotypes in normal and long-stored eggs generations F3 and F4 in female adult mosquitoes (all collection days combined). F3 mosquitoes and normal eggs storage group were blood fed. At each life stage and sex, statistical significance between genotypes was assessed with Fisher’s exact (A), Wilcoxon rank sum (B) pairwise tests. (D) Odds ratios and 95% CIs are shown for each model term from the bias-reduced logistic regression assessing factors associated with infection status. Dashed vertical line at 1 indicates no effect relative to the reference level. Asterisks indicate significance based on analysis of deviance (type III). For all panels *P ≤ 0.05, ** P ≤ 0.01, *** P ≤ 0.001, ****P ≤ 0.0001.

## Discussion

Expanding genome sequencing studies in diverse hosts have revealed that a large number of nrEVEs exists across host and virus taxa ^2,3,5,6,26–32^. However, the biological roles or evolutionary significance of these genetic elements remain poorly understood and experimental evidence demonstrating nrEVE functions *in vivo* is still scarce. In our previous work, we showed that a CFAV-derived EVE produces piRNAs and moderately restricts CFAV replication in mosquito ovaries using an artificial infection system based on intrathoracic injection ^17^. Artificial infection systems, however, do not fully recapitulate key features of persistent MSV infections, including efficient vertical transmission or long-term host-virus interaction across generations ^23,24^. A naturally infected mosquito system is particularly important because it preserves the persistent and vertically transmitted nature of MSV infections, allowing viral replication to be evaluated under physiological infection routes, tissue tropism, developmental transition, and host-virus interactions maintained across generations. This model therefore provides a more appropriate framework for determining whether nrEVEs limit viral replication in a natural context and for assessing biological relevance of such effects. To address this, we established *Ae. aegypti* mosquito lines naturally infected with CFAV that differed in the presence or absence of CFAV-EVE1, enabling us to test nrEVE-mediated antiviral effects across tissues, developmental stages, and generations under natural infection conditions.

Using this natural infection system, we demonstrate that CFAV-EVE1 suppresses replication of the cognate virus in mosquitoes via the piRNA pathway. As previously observed in CFAV-injected mosquitoes ^17^, more effective viral piRNA biogenesis was observed in the ovary, in line with higher piRNA pathway activity in this tissue.

The antiviral effect mediated by CFAV-EVE1 was observed in both female and male mosquitoes and across all life stages, from eggs to adults. Previous research on mosquito antiviral immunity has focused almost exclusively on adult females, due to their role in arbovirus transmission among humans. Whether antiviral mechanisms function broadly across sexes and developmental stages has remained largely unexplored. Our results fill this gap by demonstrating for the first time that nrEVE-mediated antiviral effects reduce viral replication across sexes and developmental stages in mosquitoes. This constitutively active, heritable antiviral system may have evolved in response to persistent, vertically transmitted viruses, thereby reducing viral burden from early life stages and allowing long-term host–virus coexistence.

To examine whether this antiviral activity is maintained across generations under natural vertical transmission, we tracked CFAV prevalence and RNA levels in CFAV-EVE1 (+/+) and (−/−) lines over nine successive generations. CFAV RNA levels were consistently lower in the CFAV-EVE1 (+/+) line, demonstrating that CFAV-EVE1 exerts a robust and stable antiviral effect. In terms of infection prevalence, the CFAV-EVE1 (−/−) line showed a gradual increase in infection prevalence, reaching fixation (100%) by the F3 generation. In contrast, the CFAV-EVE1 (+/+) line maintained a lower but persistent infection prevalence across generations, stabilizing at approximately 70–80%. The absence of a detectable difference in the F1 generation may reflect the heterozygous nature of the founding population, in which similar levels of virus would have been transmitted to F1 offspring regardless of their genotype. Consistent with this, a diverging trend of increasing prevalence in the (−/−) line and decreasing prevalence in the (+/+) line was observed from F2 onwards. The persistence of CFAV despite CFAV-EVE1-mediated suppression may be explained, at least in part, by multiple transmission routes. CFAV can be transmitted both vertically (maternal and paternal transmission) and horizontally (venereal transmission) in mosquitoes ^20^, providing multiple opportunities for infection to persist in the population even when viral replication is reduced. Differences in viral load between genotypes may further influence transmission efficiency across these routes, with higher viral loads in the CFAV-EVE1 (−/−) line potentially enhancing both vertical and venereal transmission, whereas reduced viral loads in the CFAV-EVE1 (+/+) line may limit their contribution to viral transmission. These mechanisms may help explain the observed population-level patterns. Our results indicate that CFAV-EVE1 does not eliminate infection from the population, but instead moderates viral replication and thereby tempers the long-term spread of CFAV within the host population.

Despite the clear antiviral effect, no measurable increases in fecundity, fertility, or lifespan were observed under our experimental conditions. These observations suggest that the antiviral effect of CFAV-EVE1 does not necessarily translate into detectable fitness gains in naturally infected mosquitoes when they are reared under conventional laboratory conditions. Mosquito antiviral responses often reduce viral replication without eliminating infection, and such effects may not necessarily translate into detectable fitness differences under laboratory conditions ^32^. The fitness consequences of nrEVEs may therefore be context dependent. For instance, environmental conditions are known to influence mosquito–virus interactions, and factors such as temperature, humidity, and nutritional conditions can alter viral replication and host immune responses, potentially affecting host fitness ^34–36^. Consequently, even modest EVE-mediated reductions in viral replication could become advantageous under real-world environmental conditions that are not captured in standard laboratory settings.

Consistent with this idea, our observation of a lower CFAV prevalence in mosquitoes hatched from aged eggs, particularly in individuals carrying CFAV-EVE1, suggests that nrEVEs may influence viral persistence under ecologically relevant conditions. In *Aedes* mosquitoes, eggs can remain in a prolonged quiescent state during dry periods until larval breeding sites are reflooded. Extended egg quiescence is known to alter metabolic reserves and physiological conditions in *Ae. aegypti* eggs ^37^. Vertically transmitted mosquito viruses may also persist through dry periods via ovarian infection and transovarial transmission ^38^. In this context, nrEVE-mediated suppression of viral replication could influence viral persistence across generations under ecologically relevant seasonal conditions. If such effects provide even modest advantages under natural ecological conditions, nrEVEs might be expected to be maintained by selection in mosquito populations. Recent studies have identified several CFAV-derived nrEVEs in *Ae. aegypti* genomes from geographically distinct continents ^17,25^. Population genomic analyses further indicate that CFAV-derived nrEVEs represent evolutionarily dynamic components of mosquito genomes, with some elements varying in frequency among natural populations ^25^. These observations raise the possibility that certain nrEVEs may still be shaped by selection under specific ecological conditions, potentially by providing benefits to mosquitoes in natural populations.

Alternatively, some nrEVEs may represent evolutionary relics of past viral epidemics during which strong selective pressures favored their integration and retention in host genomes. Such integrations may have contributed to long-term host–virus coevolution. Recent virome studies have revealed extensive diversity among MSVs ^39,40^, and many extant viral sequences are genetically distant from nrEVEs in mosquito genomes ^7,41^, suggesting that numerous nrEVEs may derive from extinct viruses. Over long evolutionary timescales, interactions between mosquitoes and vertically transmitted viruses may have favored the persistence of viruses with reduced pathogenicity in their hosts ^42,43^ As a result, the fitness consequences of nrEVEs may be difficult to detect under present-day laboratory conditions. At the same time, if nrEVE-derived piRNAs act as selective pressures shaping viral evolution, such effects may still be detectable in currently circulating host–virus combinations. Naturally infected mosquito lines provide an opportunity to examine nrEVE–virus interactions across multiple generations, including the potential for viral escape and host–virus arms race.

Future studies will need to consider these potential impacts to fully elucidate nrEVE-mediated antiviral functions *in vivo*. Notably, the pathogenesis of MSVs, including CFAV, remains poorly characterized in mosquitoes, particularly in the context of natural infections. Although a previous study showed that artificial infection of *Ae. aegypti* with CFAV did not result in detectable fitness costs in terms of adult survival, host-seeking behavior, or reproduction ^44^, it remains unclear whether natural, persistent CFAV infection imposes fitness costs under environmentally relevant conditions. Addressing this question will require examining whether persistent infection causes pathogenesis under environmental stress. Another key open question is whether nrEVE-associated piRNA responses represent a broadly conserved form of sequence-guided antiviral defense across animal taxa. A similar possibility has been explored in mammals using Borna disease virus 1 and endogenous bornavirus-like elements that generate piRNAs, although no detectable antiviral effect was observed, possibly due to sequence divergence between the EVE and the infecting virus or other system-specific constraints ^45^. Further studies across diverse animal models will be required to assess the generality of this mechanism.

Overall, our findings advance the functional understanding of nrEVEs by reframing them not as agents of viral elimination, but as heritable regulators of persistent virus–host interactions across generations. By limiting viral replication and transmission without eliminating infection, nrEVEs may contribute to stable long-term coexistence between hosts and their viromes. These results highlight the importance of considering persistent viruses and their endogenous genetic remnants as integral components of host biology and underscore the need to investigate nrEVE function under ecologically relevant conditions.

## Methods

### Mosquito origin and rearing

We used three laboratory colonies of *Ae. aegypti* mosquitoes: a wild-type strain derived from a natural population in Long An, Vietnam; a line homozygous for a CRISPR/Cas9-mediated knockout of CFAV-EVE1 (CFAV-EVE1 KO); and its wild-type sister line homozygous for CFAV-EVE1 ^17^. Mosquitoes were reared under controlled conditions in an environmental chamber at 28 °C, 80% relative humidity, and a 12:12 h light:dark cycle. Larvae were fed ground fish food, and adults were provided continuous access to a 10% sucrose solution.

### Detection of CFAV-EVEs by PCR

The presence of CFAV-EVEs in *Ae. aegypti* mosquito colonies was assessed by PCR. Five CFAV-EVEs previously described in the literature were targeted. The EVE and flanking sequences are shown in Table S15 ^17,25^. Total DNA was extracted from adult mosquitoes using the NucleoSpin Tissue kit (Macherey-Nagel), following the manufacturer’s instructions. PCR amplification was performed using DreamTaq DNA Polymerase (Thermo Fisher Scientific) with primer pairs listed in Table S16.

### Generation of naturally CFAV-infected *Ae. aegypti* without CFAV-EVEs

An *Ae. aegypti* line naturally infected with CFAV at nearly 100% prevalence but lacking CFAV-EVE sequences was initially established. This was achieved through multiple sequential crosses between a wild-type colony from Long An, Vietnam, and the CFAV-EVE1 KO line. Forty females from the Vietnam strain and forty males from the CFAV-EVE1 KO line were placed together in a cage and allowed to mate and produce eggs after blood feeding. Female progeny from this cross were subsequently mated with males of the CFAV-EVE1 KO line, and this crossing process was repeated two additional times. After the fourth cross, female progeny were screened for CFAV infection and for the absence of CFAV-EVE1 and CFAV-EVE2 using RT-qPCR and conventional PCR, respectively. Individuals that were positive for CFAV but negative for CFAV-EVE1 and CFAV-EVE2 were selected and maintained as a naturally CFAV-infected, EVE-deficient colony.

Second, females from the newly generated naturally CFAV-infected line lacking CFAV-EVEs were crossed with males from the sister line of the KO line retaining CFAV-EVE1. The progeny heterozygous for CFAV-EVE1 were maintained for one generation. The next generation theoretically contained three CFAV-EVE1 genotypes: (+/+), (+/−), and (−/−). To separate these genotypes, genotyping was performed using two legs from each mosquito with primers targeting the flanking region of CFAV-EVE1, as previously described ^17^, allowing individuals to be identified while remaining alive. 98 females and 35 males of the CFAV-EVE1 (+/+) or (−/−) genotypes were placed together in separate cages and maintained as naturally CFAV-infected CFAV-EVE1 (+/+) and (−/−) lines. From the F3 generation onward, each genotype was split into three replicate lines.

### CFAV RNA quantification

Total RNA was extracted from individual mosquitoes or tissues using the NucleoSpin RNA kit (Macherey-Nagel). To quantify CFAV RNA levels, RT-qPCR was performed using the PrimeScript RT reagent Kit (Perfect Real Time; Takara) using random hexamers and iTaq Universal SYBR Green Supermix (Bio-Rad). Relative CFAV RNA copy numbers were calculated by normalization to the housekeeping gene *rp49*. The primer sequences are listed in Table S1. CFAV RNA levels across generations were examined at the adult stage. To assess CFAV RNA levels during development, individuals at each stage (egg, larva, pupa, adult) were sampled and processed for quantification of CFAV RNA. Eggs and first-instar larvae were analyzed as pools of five individuals because of their small body size whereas individual specimens were used from the third-instar larval stage onward.

### RNA sequencing for virus genome recovery

Vietnam × CFAV-EVE1/2 (−/−) 4^th^ cross eggs were hatched, and fourteen individual larvae were collected on day 4, frozen at -80°C and later thawed and homogenized at 6000 rpm for 30 sec using Precellys 24 Tissue Homogenizer (Bertin Technologies) in 2 mL tubes with glass beads and 400 μL of sterile 4°C PBS, pulse-centrifuged at 4°C, and immediately chilled on ice after homogenization. 200 μL aliquot was mixed with 200 μL RNA extraction lysis buffer. 100 uL of water from the tray as well as the sterile PBS by itself were included as controls and processed in the same batch with larvae samples. Nucleic acid extraction was performed with the NucleoMag Pathogen kit (Macherey-Nagel) using IDEAL™ 32 extraction robot (ID Solutions). Extracted RNA was treated with Turbo DNase (Ambion) followed by purification using SPRI beads (Agencourt RNA clean XP, Beckman Coulter). We used a rRNA depletion approach based on RNAse H and a custom mix of oligonucleotide probes designed to target mosquito rRNA ^46^. The RNA was then subjected to randomly primed cDNA synthesis and sequencing library preparation with Nextera XT DNA Library Preparation Kit (Illumina). Libraries were sequenced on an Illumina NextSeq500 (2 × 75 cycles). Raw reads were subjected to quality control by *FastQC* v0.11.9 (https://www.bioinformatics.babraham.ac.uk/projects/fastqc/). Reads were quality-trimmed and size-filtered using *Trimmomatic* v0.39 ^47^. We mapped reads to a CFAV reference sequence from Thailand (MK860761) with *Bowtie2* v2.3.5.1 --sensitive-local alignment mode with both paired and unpaired reads to reconstruct CFAV consensus genome sequences ^48^. Aligned reads were filtered for mapping quality (-q 30) and sorted using *Samtools* v1.10 ^49^. An initial consensus sequence was generated with *iVar* v1.4.4 ^50^ using bases with quality ≥20, a minimum coverage depth of 10×, and a 50% frequency threshold. Reads were then remapped to this preliminary consensus, and a second consensus was produced with a stricter minimum coverage threshold of 100×. A final mapping round was performed against the second consensus to evaluate coverage, identify single-nucleotide variants occurring at ≥3% frequency and supported by ≥100× coverage, and generate the final consensus genome. Coverage profiles were calculated using *Bedtools* v2.29.2 ^51^, coverage and variants were visualized in *R* 4.2.3 ^52^. All final consensus sequences were submitted to GenBank (PZ436600-PZ436611).

### Small RNA sequencing and analysis

Small RNA libraries were prepared by Novogene using NEBNext® Multiplex Small RNA Library Prep Set for Illumina® (New England Biolabs, Cat. E7300) and sequenced on an Illumina NovaSeq 6000 short-read sequencer. After initial quality control by the sequencing facility (using in-house protocols), the obtained sequencing libraries were analyzed using the Galaxy bioinformatics tool shed (usegalaxy.org) ^53,54^. Sequencing adapters were trimmed using ‘Clip adapter sequence’(Galaxy Version 1.0.3+galaxy2) with default settings and the adapter sequence AGATCGGAAGAGCACACGTCTGAACTCCAGTCAC. Quality control of the libraries was then performed using FastQC (Galaxy Version 0.74+galaxy1). Trimmed reads were mapped against the CFAV genome (ENA: LR596014) and *Ae. aegypti* pre-microRNAs sequences (downloaded from miRBase on August 26^th^ 2025) was done using Bowtie2 ^55^ (Galaxy version 2.5.4+galaxy0) with default settings and SAM output. Continuously mapped reads were used to generate small RNA size profiles and genome profiles. Read counts were normalized by the number of mapped microRNA reads in each library. For size profiles, the normalized read count for each read length was plotted. For genome profiles of piRNAs, 25-30nt sized reads were selected and the frequency of the 5’ terminal nt was determined for each nt position of the CFAV genome. For the piRNA-sized reads nucleotide biases were determined using ‘Sequence logo generator’ (Galaxy Version 3.5.0) ^56^. Small RNA overlap probabilities were determined using ‘Small RNA signatures’ (Galaxy Version 3.5.0) available through the Mississippi[2] instance of galaxy [https://mississippi.sorbonne-universite.fr]. BAM-formatted alignment files were used as input to compute overlap signatures for 25-30 nt (min-max) reads with a minimum of 1-nt and maximum of 25-nt overlap being plotted. Small RNA sequencing data is deposited to SRA (PRJNA1481434).

### Mosquito survival curves

Forty adult mosquitoes (5–7 days post-emergence) from the CFAV-EVE1 (+/+) or (−/−) line were placed in a cage, and their survival was monitored daily. To account for mosquito mortality caused by cold anesthesia, individuals that died the following day were excluded from the survival analysis. At least 30 surviving mosquitoes per line were included in the final survival analysis.

### Artificial blood-feeding, fecundity, and fertility

CFAV-EVE1 (+/+) and (−/−) lines were blood-fed on 37°C-warmed rabbit blood for 15 min using an artificial blood-feeding apparatus (Hemotek). Blood-fed females were individually isolated in fly vials containing cotton with water-moistened paper for oviposition and provided with access to a 10% sucrose solution. At day 7 post blood-feeding, the laid eggs were counted and immediately transferred to a 12-well plate containing 2 ml water. The number of hatched larvae was counted 5 days after the egg transfer.

### Statistical analyses

Statistical analyses were performed in GraphPad Prism and R v4.2.3 and v4.4.0. Within each experimental group, infection status (CFAV RNA presence) and infection levels (CFAV RNA levels) in infected mosquitoes were compared between genotypes using pairwise Wilcoxon rank-sum and Fisher’s exact tests, respectively, applying Bonferroni correction to adjust P-values for multiple comparisons. Because each genotype included three replicate mosquito lines, the same pairwise testing framework was also applied within each genotype to evaluate differences between replicate lines in each group within each CFAV-EVE1 genotype.

In generation F1, the relationship between RNA levels and genotype (CFAV-EVE1(−/−), CFAV-EVE1(–/+), CFAV-EVE1(+/+)) was analyzed in R. Non-parametric associations were verified with Spearman rank correlations, where genotype was treated as an ordered factor coded numerically as follows: (−/−) = 1, (+/–) = 2, and (+/+) = 3.

Mosquito survival data were analyzed separately in males and females using the R packages tidyverse, survival, coxme, and survminer. The Kaplan-Meier was plotted and visually inspected (Figure S6). A mixed-effects Cox proportional hazards model was fitted to assess the impact of the mosquito genotype (CFAV-EVE1 (−/−) or without CFAV-EVE1 (+/+)), while accounting for variations in replicate lines (three per genotype).

To examine broader effects on infection status, generalized linear models (logit link and binomial error distribution) were fitted in R using the tidyverse and brglm2 packages to test the effects of genotype, egg storage conditions (normal vs. long), and mosquito generation (F4 vs. F5), life stage (combined with sex into single variable: third-instar larvae, pupae, adult female day 1, adult male day 1, adult female day 7 and adult male day 7), and body part. Bias-reduced estimation (brglm2) was due to quasi-separation issues where infection was present in all individuals in some groups (in EVE (−/−)). Model selection was based on likelihood-ratio tests, analyses of deviance, and AIC comparison, with Type III tests performed using the car package. Observed infection proportions (± 95 % binomial CI) were summarized and visualized in ggplot2. For the RNA levels, infection data were log10-transformed and, only for the linear models, outliers removed based on the visual inspection of the data. Linear models or, where possible, linear mixed-effects models were then fitted with lme4, including fixed effects and replicate mosquito line nested within genotype as a random effect. Successive models excluded insignificant interactions as well as zero or near-zero random effect of the mosquito line.

Experimental data and pairwise testing results are available in Data S1-7.

## Acknowledgements

We thank Sora Omura for assistance with mosquito colony maintenance, and Anna Beth Crist for generating the CFAV-EVE1 KO line and its wild-type sister line.

## Funding

This work was supported by JSPS KAKENHI Grant Number JP21KK0107, 21K19119, 21H02206. This study was also supported by the Ministry of Education, Culture, Sports, Science and Technology, Japan (MEXT) to a project on Joint Usage/Research Center– Leading Academia in Marine and Environment Pollution Research (LaMer). YS was supported by the Ministry of Education, Culture, Sports, Science and Technology, Japan (MEXT) to a project on Promotion of the establishment of Asian research hubs by reforming the Center for Marine Environmental Studies. The E.S.-L. laboratory is funded by Institut Pasteur, the INCEPTION program (Investissements d’Avenir grant ANR-16-CONV-0005), the Ixcore foundation for research, the French Government’s Labex IBEID (ANR-10-LABX-62-IBEID), the HERA Project DURABLE (grant no 101102733) and LEAPS (grant no 101094685).

## Author Contributions

Conceptualization, Y.Su.; Formal analysis, Y.Su., A.B., P.M.; Funding acquisition, Y.Su., A.B., K.W.; Investigation, Y.Su., A.B., Y.Se., I.C.A.A., and M.P.; Methodology, Y.Su., A.B., P.M.; Project administration, Y.Su.; Resources, Y.Su., L.L., E.S.-L., K.W.; Visualization, Y.Su., A.B., P.M.; Writing – original draft, Y.Su., A.B.; Writing – review & editing, Y.Su., A.B., Y.Se., I.C.A.A., P.M., M.P., L.L., E.S.-L., K.W.

## Competing interests

The authors declare no competing interests.

## Notes

### Competing Interest Statement

The authors have declared no competing interest.

