## Supplementary material for "Non-retroviral Endogenous Viral Element Acts as a Stable Antiviral Regulator Across Mosquito Life Stages and Generations": Supplemantary figure 1-7

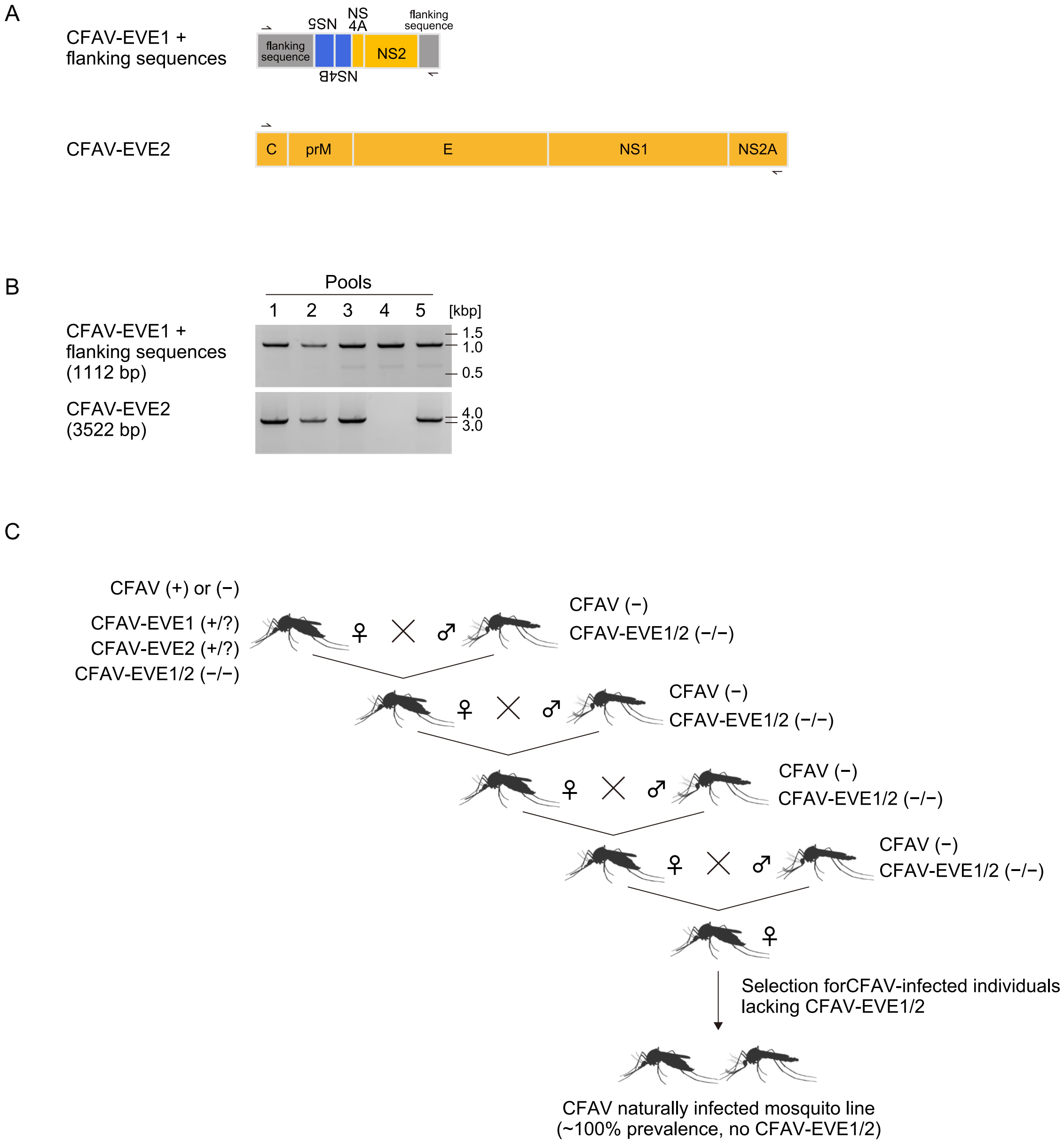


**Figure. S1. Generation of a naturally CFAV-infected mosquito line lacking CFAV-EVEs.**

(A) Schematic illustration of CFAV-EVE1 with flanking regions and CFAV-EVE2, showing primer binding sites for PCR detection. (B) PCR-based detection of CFAV-EVE1 and CFAV-EVE2 using five pools of mosquito DNA, each consisting of 5 individuals (lane 1-5).[LL8.1] (C) Schematic illustration of the crossing scheme used to establish a naturally CFAV-infected Ae. aegypti line lacking CFAV-EVEs.

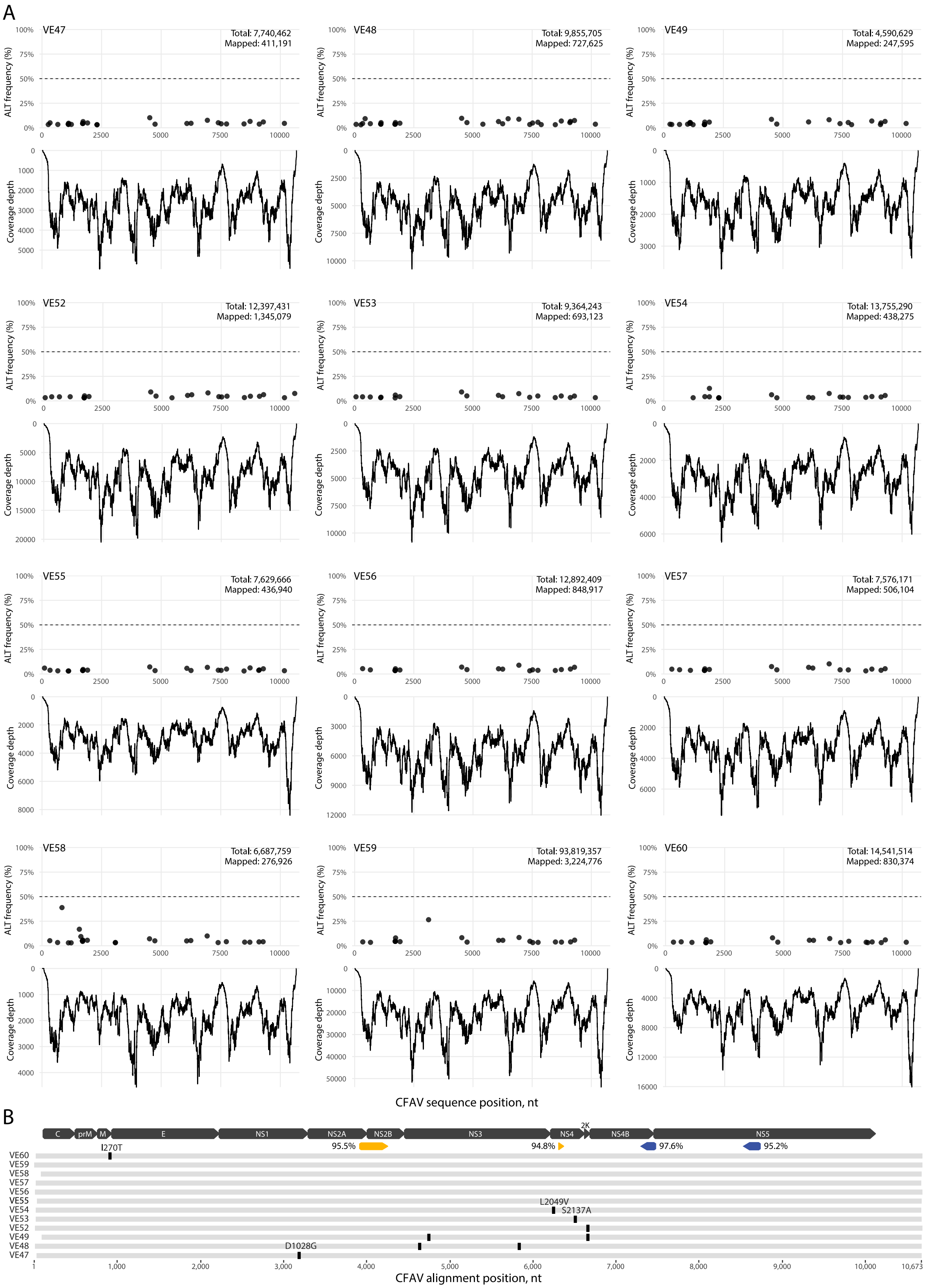


**Figure. S2. Low diversity among CFAV sequences in the parental Vietnam × CFAV-EVE1/2 (−/−) 4th**

**cross mosquitoes before the main experimental cross.**

(A) Single nucleotide variant frequencies are shown on the top sub-panels and read coverage depth on the bottom for each of the individual larvae samples (IDs indicated in the top-left corners). The total number of quality-trimmed and filtered reads as well as the number of mapped to CFAV reads are indicated in the top-right corners. CFAV position refers to the position in consensus of second mapping round. (B) Nucleotide alignment of all novel CFAV consensus sequences after the third and final read mapping round. The nucleotide differences are shown with black vertical bars, and non-synonymous ones are additionally annotated. CFAV genome map is represented above the alignment with protein names and CFAV-EVE1 fragments are colored in yellow (forward) and blue (reverse) with nucleotide identity to the novel sequences annotated.

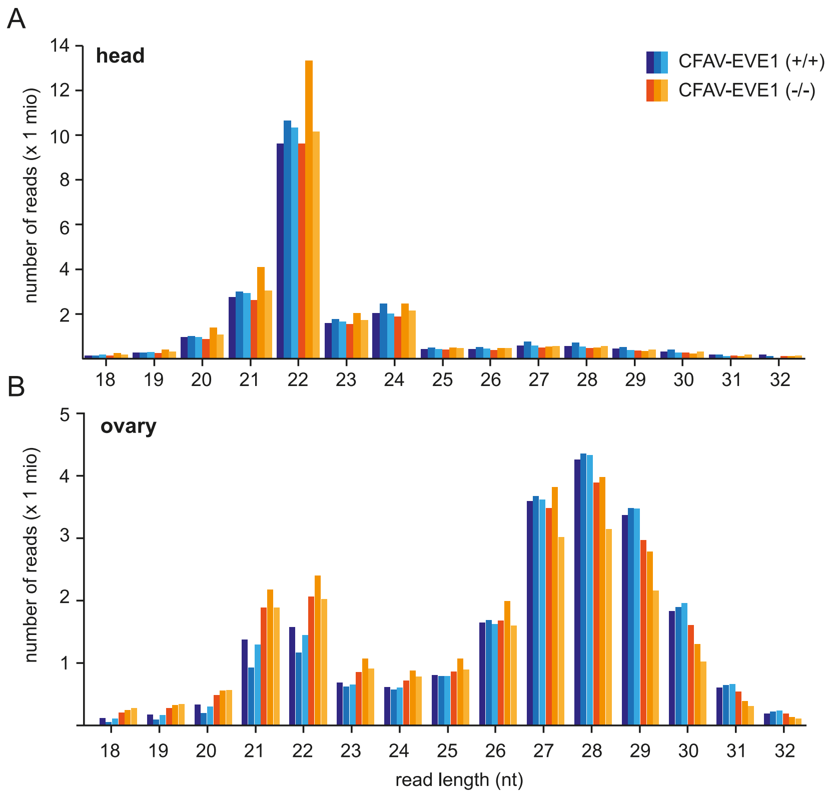


**Figure S3. Size distribution of total small RNAs from naturally CFAV-infected mosquito lines with or without CFAV-EVE1.**

Size distribution of total small RNAs from (A) head and (B) ovary samples dissected from naturally CFAV-infected Ae. aegypti with or without CFAV-EVE1. Blue and orange color gradients indicate individual replicate lines for CFAV-EVE1 (+/+) and (−/−) genotypes, respectively.

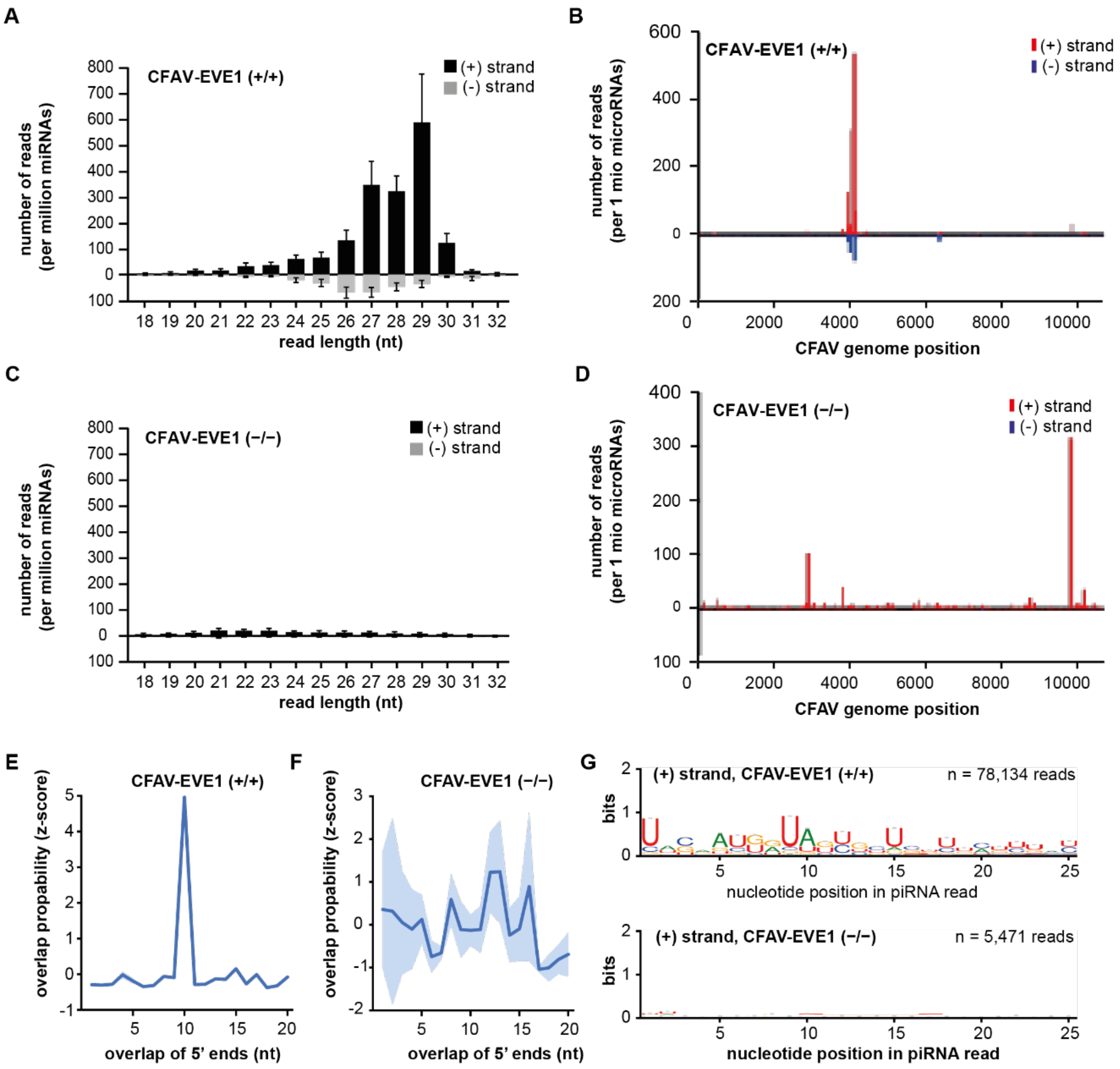


**Figure S4. CFAV-EVE1-derived piRNAs interact with CFAV RNA in naturally infected mosquito heads.**

Size distribution (A, C), genome-wide distribution of piRNA-like small RNAs (25–30 nt) (B, D), overlap probability analysis of 25–30 nt small RNAs (E, F), and sequence logo analysis depicting nucleotide bias of 25–30 nt small RNAs (G, H) mapping to the CFAV genome in heads of CFAV-EVE1 (+/+) and (−/−) mosquitoes, respectively.

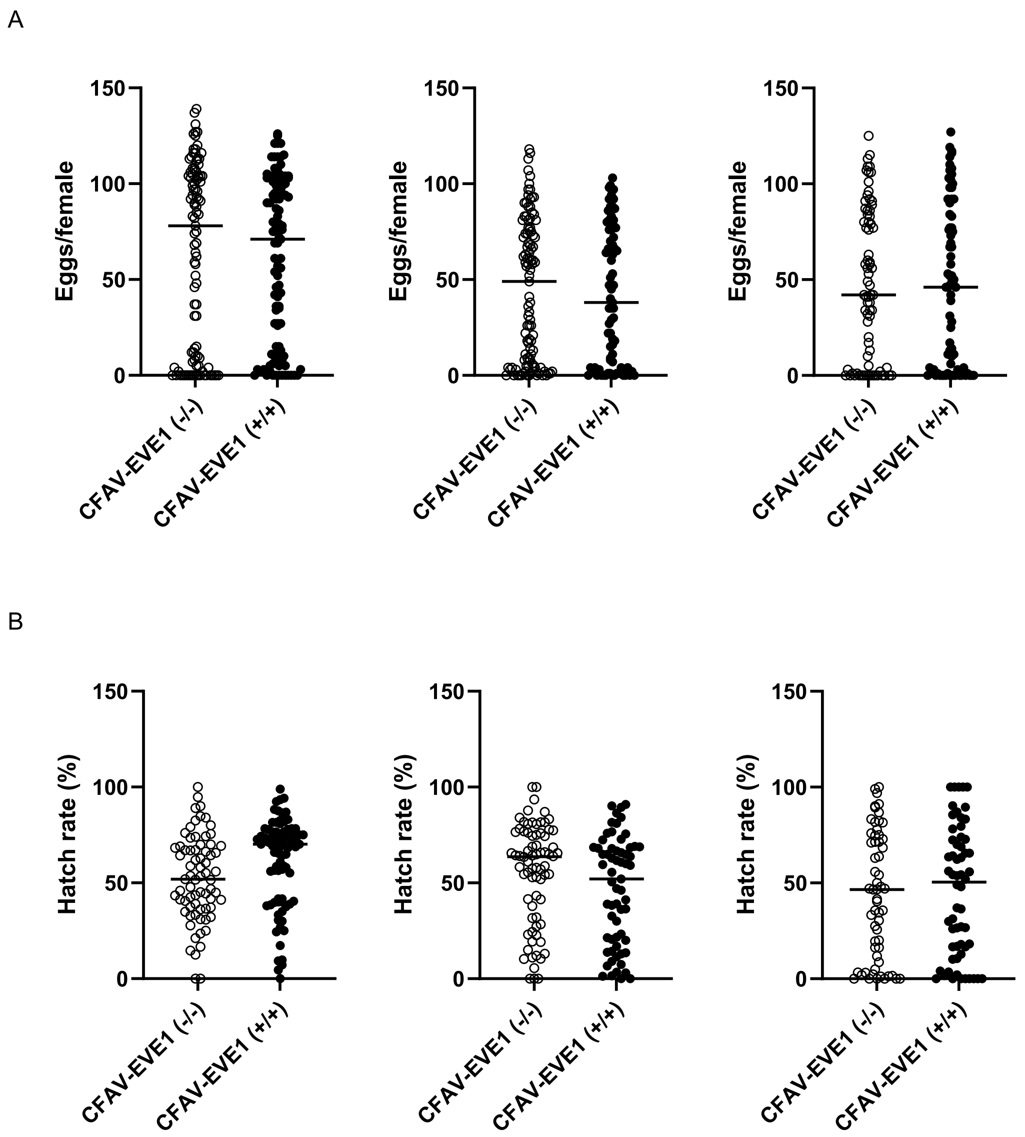


**Figure S5. CFAV-EVE1 does not affect fecundity or fertility of naturally CFAV-infected *Ae. aegypti.***

(A) Number of eggs laid and (B) proportion of eggs hatching per individual female in CFAV-EVE1 (+/+) and (−/−) mosquitoes. Bars represent the mean. Statistical significance was assessed using the Wilcoxon rank sum test.

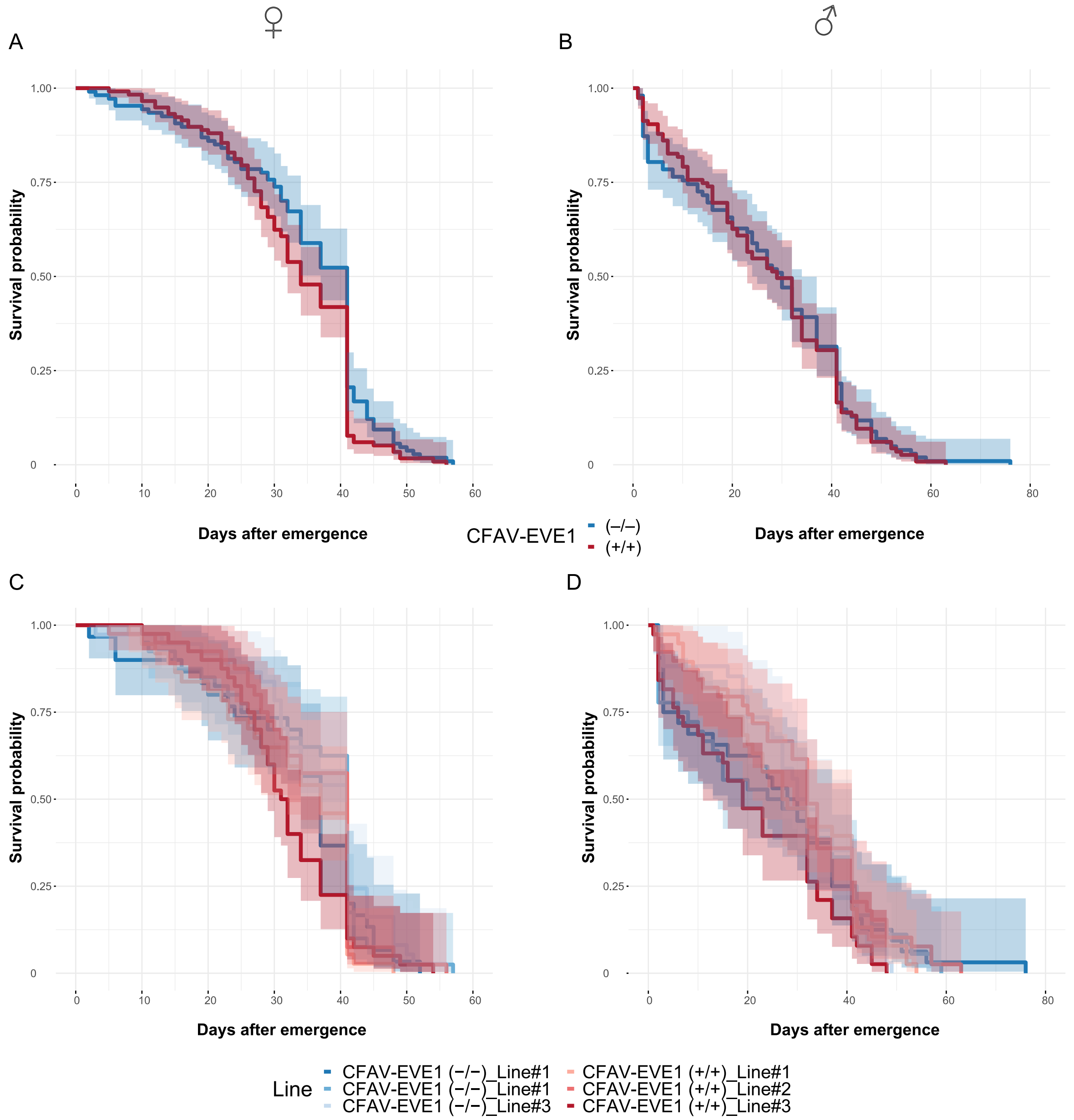


**Figure S6. Survival of naturally CFAV infected Ae. aegypti adults with or without CFAV-EVE1.**
Kaplan–Meier survival curves showing the probability of survival over time for female (A, C) or male (B, D) mosquitoes by (A, B) CFAV-EVE1 genotype ((−/−), (+/+)) or (C, D) replicate line nested within each genotype. Shaded areas represent 95 % confidence intervals around the estimated survival curves.

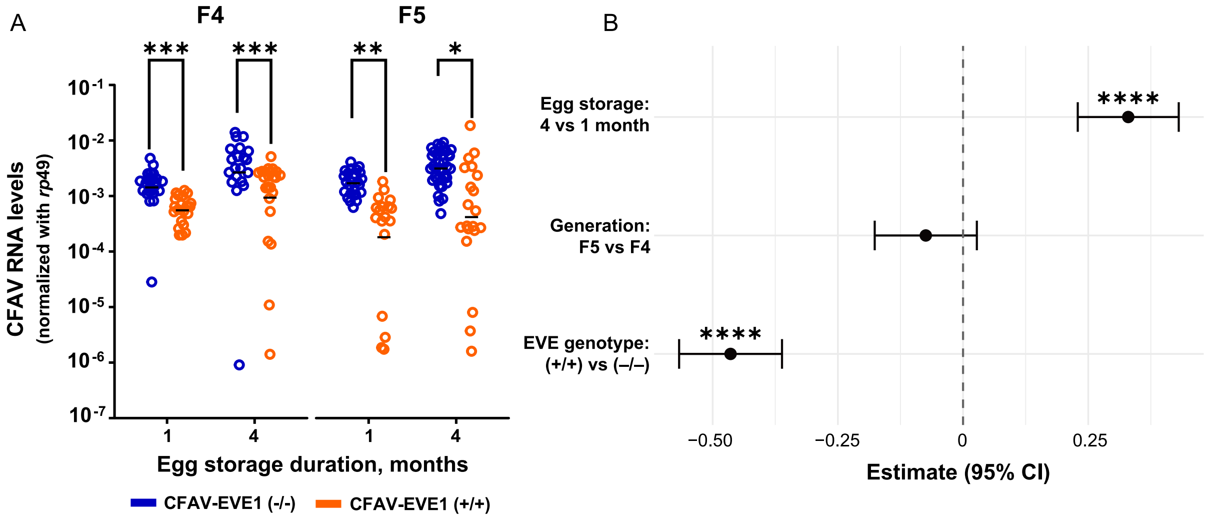


**Figure S7. Effect of prolonged egg storage on CFAV RNA level dynamics between CFAV-EVE1 genotypes.**

(A) CFAV RNA levels in mosquitoes derived from eggs subjected to normal or prolonged storage, shown for two generations. CFAV RNA levels were normalized to rp49 transcripts. Each dot represents an individual mosquito. Statistical significance between genotypes was assessed with Wilcoxon rank sum test. (B) Estimated coefficients (± 95 % CI) from the linear model on $\log_{10}$-transformed RNA levels. The outliers ≤10^-4 were removed from the dataset used for the model. Positive values denote higher mean infection compared with the reference category, negative values denote lower infection. Significance based on White-adjusted ANOVA (type III) is shown. For both panels $^{*}P \leq 0.05$, $^{**}P \leq 0.01$, $^{***}P \leq 0.001$, $^{****}P \leq 0.0001$.
