## Supplementary table 1-15 for "Non-retroviral Endogenous Viral Element Acts as a Stable Antiviral Regulator Across Mosquito Life Stages and Generations"

<table>
<tr><td colspan="3">Model: logit [Infection_status ~ Geno_num]</td><td></td><td></td><td></td></tr>
<tr><td>Coefficients:</td><td>Estimate</td><td>Std. Error</td><td>z value</td><td>Pr(>|z|)</td><td></td></tr>
<tr><td>(Intercept)</td><td>6.320</td><td>2.822</td><td>2.239</td><td>0.0251</td><td>*</td></tr>
<tr><td>Geno_num</td><td>-1.636</td><td>1.021</td><td>-1.603</td><td>0.1090</td><td></td></tr>
<tr><td colspan="3">Null deviance: 27.360 on 46 degrees of freedom</td><td></td><td></td><td></td></tr>
<tr><td colspan="3">Residual deviance: 23.305 on 45 degrees of freedom</td><td></td><td></td><td></td></tr>
<tr><td>AIC: 27.305</td><td></td><td></td><td></td><td></td><td></td></tr>
<tr><td colspan="6">Type of estimator: AS_mixed (mixed bias-reducing adjusted score equations)</td></tr>
<tr><td colspan="6">Number of Fisher Scoring iterations: 6</td></tr>
<tr><td></td><td></td><td></td><td></td><td></td><td></td></tr>
<tr><td>Analysis of Deviance</td><td>LR Chisq</td><td>Df</td><td colspan="2">Pr(>Chisq)</td><td></td></tr>
<tr><td>Geno_num</td><td>4.055</td><td>1</td><td>0.04404</td><td>*</td><td></td></tr>
</table>

**Table S1.**

Logistic model of CFAV infection status as a function of CFAV-EVE1 genotype (−/−, +/−, +/+ substituted with numbers 1, 2, 3, respectively) in males only of F1 generation (Exp02). The separately tested effect of genotype in females was not significant. Related to Figure 1B.

| Experiment | Generation | Sex | ρ (Spearman's rho) | S statistic | p-value | |
|---|---|---|---|---|---|---|
| Exp2 | F2 | Female | −0.558 | 17882 | 0.0001517 | **** |
| Exp2 | F2 | Male | −0.623 | 21500 | 8.005e-06 | **** |

**Table S2.**

Spearman's correlation of CFAV-EVE1 genotype (−/−, +/−, +/+ substituted with numbers 1, 2, 3, respectively) with $\log_{10}$-transformed CFAV RNA levels. Related to Figure 1B.

| | | | | | |
|---|---|---|---|---|---|
| Model: $\log_{10}$ Infection ~ Genotype × Body part | | | | | |
| Coefficients | Estimate | Std. Error | t value | Pr(>\|t\|) | |
| Intercept | -2.35492 | 0.1282 | -18.37 | <2.00e-16 | **** |
| Genotype++ | -0.37607 | 0.17093 | -2.2 | 0.0333 | * |
| Body_partHead | 0.18623 | 0.1813 | 1.027 | 0.3102 | |
| Body_partOvary | -1.69225 | 0.1813 | -9.334 | 8.42e-12 | **** |
| Genotype++:Body_partHead | -0.27769 | 0.24173 | -1.149 | 0.2572 | |
| Genotype++:Body_partOvary | -0.05517 | 0.24173 | -0.228 | 0.8206 | |
| Residual standard error: 0.3392 on 42 degrees of freedom | | | | | |
| Multiple R-squared: 0.8793, Adjusted R-squared: 0.865 | | | | | |
| F-statistic: 61.21 on 5 and 42 DF, p-value: < 2.2e-16 | | | | | |
| Test of model assumptions | Statistic | p-value | | | |
| Shapiro–Wilk normality | W = 0.941 | 0.018 | * | | |
| Levene's homogeneity | F(5, 42) = 2.21 | 0.071 | | | |
| Type III ANOVA | Sum Sq | Df | F value | p-value | |
| Intercept | 38.82 | 1 | 337.44 | <2.00e-16 | **** |
| Genotype | 0.557 | 1 | 4.8408 | 0.03335 | * |
| Body_part | 14.997 | 2 | 65.18 | 1.33e-13 | **** |
| Genotype:Body_part | 0.17 | 2 | 0.7397 | 0.48337 | |
| Residuals | 4.832 | 42 | | | |

**Table S3.**

Linear model and ANOVA summary of CFAV-EVE1 genotype and body part effects on $\log_{10}$-transformed CFAV RNA infection levels (Females; Generation F4, Exp03). Related to Figure 1C.

| Cox mixed-effects model: surv_obj_dff ~ EVE + (1 \| Line) | | | | | |
|---|---|---|---|---|---|
| events n = 217 | | | | | |
| Random effects: | | | | | |
| | group | variable | sd | variance | |
| 1 | Line | Intercept | 0.112709 | 1.27E-02 | |
| | Chisq | df | p | AIC | BIC |
| Integrated loglik | 0.54 | 2 | 0.764 | -3.46 | -10.2 |
| Penalized loglik | 3.94 | 2.23 | 0.1664 | -0.52 | -8.05 |
| Fixed effects: | | | | | |
| | coef | exp(coef) | se(coef) | z | p |
| EVEpresent | 0.077 | 1.08 | 0.1651 | 0.47 | 0.641 |
| PH-assumption check (clustered by Line; using cox.zph() function) | | | | | |
| | chisq | df | p | | |
| EVE | 0.444 | 1 | 0.5 | | |
| GLOBAL | 0.444 | 1 | 0.5 | | |

**Table S4.**

Cox mixed-effects model for survival analysis in males, generation F4, Exp19. Related to Figure S6B.

| | | | | | | |
|---|---|---|---|---|---|---|
| Cox mixed-effects model: surv_obj_dff ~ EVE + (1 \| Line) | | | | | | |
| events n = 224 | | | | | | |
| Random effects: | | | | | | |
| | group | variable | sd | variance | | |
| 1 | Line | Intercept | 0.00915 | 8.37E-05 | | |
| | Chisq | df | p | AIC | BIC | |
| Integrated loglik | 5.15 | 2 | 0.07627 | 1.15 | -5.68 | |
| Penalized loglik | 5.17 | 1.01 | 0.02339 | 3.15 | -0.3 | |
| Fixed effects: | | | | | | |
| | coef | exp(coef) | se(coef) | z | p | |
| EVEpresent | 0.3073 | 1.3597 | 0.1357 | 2.26 | 0.024 | * |
| PH-assumption check (clustered by Line; using cox.zph() function) | | | | | | |
| | chisq | df | p | | | |
| EVE | 0.131 | 1 | 0.72 | | | |
| GLOBAL | 0.131 | 1 | 0.72 | | | |

**Table S5.**

Cox mixed-effects model for survival analysis in females, generation F4, Exp19. Related to Figure S6A.

| Model: logit [Infection_status ~ Genotype × Generation] | | | | | |
|---|---|---|---|---|---|
| Coefficients: | Estimate | Std. Error | z value | Pr(>\|z\|) | |
| Intercept | 1.3545 | 0.3966 | 3.416 | 0.000636 | *** |
| Genotype++ | 0.5914 | 0.6212 | 0.952 | 0.341074 | |
| GenerationF2 | 1.0433 | 0.8383 | 1.245 | 0.213267 | |
| Genotype++:GenerationF2 | -2.2961 | 1.0577 | -2.171 | 0.029946 | * |
| Null deviance: 120.16 on 126 degrees of freedom | | | | | |
| Residual deviance: 114.04 on 123 degrees of freedom | | | | | |
| AIC: 122.04 | | | | | |
| Number of Fisher Scoring iterations: 5 | | | | | |
| Analysis of Deviance Type III | LR Chisq | Df | Pr(>Chisq) | | |
| Genotype | 0.9285 | 1 | 0.3353 | | |
| Generation | 1.7847 | 1 | 0.1816 | | |
| Genotype:Generation | 5.3054 | 1 | 0.0213 | * | |

**Table S6.**

Logistic model of CFAV infection rate as a function of CFAV-EVE1 genotype and generation (females; Generations F2–F3, pooled Exp02 and Exp15 for F2 (no statistically significant difference between experiments within F2), Exp14 for F3). Related to Figure 3A.

| Model: logit [Infection_status ~ Genotype + Generation] | | | | | |
|---|---|---|---|---|---|
| Coefficients: | Estimate | Std. Error | z value | Pr(>\|z\|) | |
| (Intercept) | 2.4103 | 0.6709 | 3.592 | 0.000328 | *** |
| Genotype++ | -1.8097 | 0.7447 | -2.43 | 0.015093 | * |
| GenerationF4 | 1.3588 | 0.6783 | 2.003 | 0.045157 | * |
| Null deviance: 76.150 on 95 degrees of freedom | | | | | |
| Residual deviance: 63.547 on 93 degrees of freedom | | | | | |
| AIC: 69.547 | | | | | |
| Type of estimator: AS_mixed (mixed bias-reducing adjusted score equations) | | | | | |
| Number of Fisher Scoring iterations: 3 | | | | | |
| Analysis of Deviance Type II | LR Chisq | Df | Pr(>Chisq) | | |
| Genotype | 8.0938 | 1 | 0.0044 | ** | |
| Generation | 4.8544 | 1 | 0.0276 | * | |

**Table S7.**

Bias-reduced logistic model of CFAV infection status as a function of CFAV-EVE1 genotype and generation F3 and F4 (females; F3 in Exp14, F4 in Exp04). Related to Figure 3A.

| Model: logit [Infection_status ~ Genotype + Generation] | | | | | |
|---|---|---|---|---|---|
| Coefficients: | Estimate | Std. Error | z value | Pr(>\|z\|) | |
| (Intercept) | 6.8258 | 1.4867 | 4.591 | 4.41E-06 | **** |
| Genotype++ | -5.0103 | 1.3788 | -3.634 | 0.000279 | *** |
| GenerationF5 | -0.7692 | 0.7463 | -1.031 | 0.302687 | |
| GenerationF6 | -0.9679 | 0.7341 | -1.319 | 0.187327 | |
| GenerationF7 | -1.1518 | 0.7253 | -1.588 | 0.112254 | |
| GenerationF8 | -1.1518 | 0.7253 | -1.588 | 0.112254 | |
| GenerationF9 | -0.8055 | 0.6428 | -1.253 | 0.210146 | |
| GenerationF10 | -0.5496 | 0.7637 | -0.72 | 0.471754 | |
| Null deviance: 333.23 on 431 degrees of freedom | | | | | |
| Residual deviance: 243.99 on 424 degrees of freedom | | | | | |
| AIC: 259.99 | | | | | |
| Type of estimator: AS_mixed (mixed bias-reducing adjusted score equations) | | | | | |
| Number of Fisher Scoring iterations: 7 | | | | | |
| Analysis of Deviance Type II | LR Chisq | Df | Pr(>Chisq) | | |
| Genotype | 85.605 | 1 | <2e-16 | **** | |
| Generation | 4.234 | 6 | 0.6451 | | |

**Table S8.**

Bias-reduced logistic model of CFAV infection status as a function of CFAV-EVE1 genotype and generation F4–F10 (females; F4 – Exp04, F5–F10 from Exp07–Exp13). Related to Figure 3A.

| Model: logit [Infection_status ~ Genotype + Generation] | | | | | |
|---|---|---|---|---|---|
| Coefficients: | Estimate | Std. Error | z value | Pr(>\|z\|) | |
| (Intercept) | 6.04901 | 1.45251 | 4.165 | 3.12E-05 | **** |
| Genotype++ | -4.53217 | 1.37499 | -3.296 | 0.00098 | *** |
| GenerationF6 | -0.25058 | 0.71879 | -0.349 | 0.72738 | |
| GenerationF7 | -0.25058 | 0.71879 | -0.349 | 0.72738 | |
| GenerationF8 | -0.25058 | 0.71879 | -0.349 | 0.72738 | |
| GenerationF9 | -0.03251 | 0.60721 | -0.054 | 0.9573 | |
| GenerationF10 | -0.47015 | 0.70034 | -0.671 | 0.50201 | |
| Null deviance: 247.91 on 383 degrees of freedom | | | | | |
| Residual deviance: 191.50 on 377 degrees of freedom | | | | | |
| AIC: 205.5 | | | | | |
| Type of estimator: AS_mixed (mixed bias-reducing adjusted score equations) | | | | | |
| Number of Fisher Scoring iterations: 7 | | | | | |
| Analysis of Deviance Type II | LR Chisq | Df | Pr(>Chisq) | | |
| Genotype | 55.934 | 1 | 7.50e-14 | **** | |
| Generation | 0.54 | 5 | 0.9906 | | |

**Table S9.**

Bias-reduced logistic model of CFAV infection status as a function of CFAV-EVE1 genotype and generation F5–F10 (males; F5–F10 from Exp07–Exp13). Related to Figure 3B.

<table>
<tr><td colspan="4">Model: $\log_{10}$ Infection ~ Genotype + Generation + (1 | Line)</td><td></td><td></td><td></td></tr>
<tr><td>Random effects:</td><td></td><td></td><td></td><td></td><td></td><td></td></tr>
<tr><td>Groups</td><td>Name</td><td>Variance</td><td>Std.Dev.</td><td></td><td></td><td></td></tr>
<tr><td>Line</td><td>(Intercept)</td><td>0.005324</td><td>0.07296</td><td></td><td></td><td></td></tr>
<tr><td>Residual</td><td></td><td>0.121809</td><td>0.34901</td><td></td><td></td><td></td></tr>
<tr><td>Fixed effects:</td><td>Estimate</td><td>Std. Error</td><td>df</td><td>t value</td><td>Pr(>|t|)</td><td></td></tr>
<tr><td>(Intercept)</td><td>-3.09318</td><td>0.07211</td><td>19.32266</td><td>-42.897</td><td>< 2e-16</td><td>****</td></tr>
<tr><td>Genotype++</td><td>-0.50371</td><td>0.06913</td><td>4.14536</td><td>-7.286</td><td>0.001641</td><td>**</td></tr>
<tr><td>GenerationF4</td><td>0.29033</td><td>0.07692</td><td>395.18474</td><td>3.775</td><td>0.000185</td><td>***</td></tr>
<tr><td>GenerationF5</td><td>0.33504</td><td>0.08023</td><td>396.09319</td><td>4.176</td><td>3.65E-05</td><td>****</td></tr>
<tr><td>GenerationF6</td><td>0.48843</td><td>0.07908</td><td>395.26486</td><td>6.176</td><td>1.63E-09</td><td>****</td></tr>
<tr><td>GenerationF7</td><td>0.47796</td><td>0.07911</td><td>395.4214</td><td>6.042</td><td>3.52E-09</td><td>****</td></tr>
<tr><td>GenerationF8</td><td>0.31865</td><td>0.07957</td><td>395.18214</td><td>4.004</td><td>7.43E-05</td><td>****</td></tr>
<tr><td>GenerationF9</td><td>0.0759</td><td>0.06467</td><td>395.28168</td><td>1.174</td><td>0.241272</td><td></td></tr>
<tr><td>GenerationF10</td><td>-0.01142</td><td>0.07773</td><td>395.23986</td><td>-0.147</td><td>0.883235</td><td></td></tr>
<tr><td></td><td></td><td></td><td></td><td></td><td></td><td></td></tr>
<tr><td colspan="7">Analysis of Deviance Table (Type II Wald chisquare tests)</td></tr>
<tr><td></td><td>Chisq</td><td>Df</td><td>Pr(>Chisq)</td><td></td><td></td><td></td></tr>
<tr><td>(Intercept)</td><td>1840.114</td><td>1</td><td>< 2e-16</td><td>****</td><td></td><td></td></tr>
<tr><td>Genotype</td><td>53.088</td><td>1</td><td>3.19E-13</td><td>***</td><td></td><td></td></tr>
<tr><td>Generation</td><td>109.464</td><td>7</td><td>< 2e-16</td><td>****</td><td></td><td></td></tr>
</table>

**Table S10.**

Linear mixed-effects model of $\log_{10}$-transformed CFAV infection levels as a function of CFAV-EVE1 genotype and generation F3–F10 (females; F3 – Exp14, F4– Exp04, F5–F10 from Exp07–Exp13, respectively). Outliers <$10^{-5}$ were removed from the dataset. Related to Figure 3C.

<table>
<tr><td colspan="4">Model $\log_{10}$ Infection ~ Genotype + Generation + (1 | Line)</td><td></td><td></td><td></td></tr>
<tr><td>Random effects:</td><td></td><td></td><td></td><td></td><td></td><td></td></tr>
<tr><td>Groups</td><td>Name</td><td>Variance</td><td>Std.Dev.</td><td></td><td></td><td></td></tr>
<tr><td>Line</td><td>(Intercept)</td><td>0.001514</td><td>0.03891</td><td></td><td></td><td></td></tr>
<tr><td>Residual</td><td></td><td>0.076087</td><td>0.27584</td><td></td><td></td><td></td></tr>
<tr><td>Fixed effects:</td><td>Estimate</td><td>Std. Error</td><td>df</td><td>t value</td><td>Pr(>|t|)</td><td></td></tr>
<tr><td>(Intercept)</td><td>-1.83754</td><td>0.05034</td><td>25.8391</td><td>-36.5</td><td>< 2e-16</td><td>****</td></tr>
<tr><td>Genotype++</td><td>-0.49465</td><td>0.04369</td><td>3.77341</td><td>-11.323</td><td>0.000475</td><td>***</td></tr>
<tr><td>GenerationF6</td><td>0.07623</td><td>0.06024</td><td>332.41911</td><td>1.265</td><td>0.206633</td><td></td></tr>
<tr><td>GenerationF7</td><td>0.04899</td><td>0.06021</td><td>331.83043</td><td>0.814</td><td>4.16E-01</td><td></td></tr>
<tr><td>GenerationF8</td><td>-0.05089</td><td>0.06023</td><td>332.15268</td><td>-0.845</td><td>3.99E-01</td><td></td></tr>
<tr><td>GenerationF9</td><td>-0.31875</td><td>0.04938</td><td>332.07869</td><td>-6.456</td><td>3.82E-10</td><td>****</td></tr>
<tr><td>GenerationF10</td><td>0.05205</td><td>0.06059</td><td>332.40462</td><td>0.859</td><td>3.91E-01</td><td></td></tr>
<tr><td></td><td></td><td></td><td></td><td></td><td></td><td></td></tr>
<tr><td colspan="7">Analysis of Deviance Table (Type II Wald chisquare tests)</td></tr>
<tr><td></td><td>Chisq</td><td>Df</td><td>Pr(>Chisq)</td><td></td><td></td><td></td></tr>
<tr><td>(Intercept)</td><td>1332.25</td><td>1</td><td>< 2e-16</td><td>****</td><td></td><td></td></tr>
<tr><td>Genotype</td><td>128.2</td><td>1</td><td>< 2e-16</td><td>****</td><td></td><td></td></tr>
</table>

**Table S11.**

Linear mixed-effects model of $\log_{10}$-transformed CFAV infection levels as a function of CFAV-EVE1 genotype and generation F3–F10 (males; from Exp07–Exp13, respectively). Outliers $<10^{-4}$ were removed from the dataset. Related to Figure 3D.

| Model: logit [Infection_status ~ Genotype + Stage_Sex] | | | | | |
|---|---|---|---|---|---|
| Coefficients: | Estimate | Std. Error | z value | Pr(>\|z\|) | |
| (Intercept) | 4.7707 | 1.3872 | 3.439 | 0.000584 | *** |
| Genotype++ | -4.2786 | 1.371 | -3.121 | 0.001804 | ** |
| Stage_SexPupa | 1.0251 | 0.6694 | 1.531 | 0.125661 | |
| Stage_SexAdult_D1_Female | 0.7746 | 0.6394 | 1.212 | 0.22568 | |
| Stage_SexAdult_D1_Male | 1.3239 | 0.7151 | 1.851 | 0.064098 | . |
| Stage_SexAdult_D7_Female | 1.0251 | 0.6694 | 1.531 | 0.125661 | |
| Stage_SexAdult_D7_Male | 1.3239 | 0.7151 | 1.851 | 0.064098 | . |
| Null deviance: 183.71 on 287 degrees of freedom | | | | | |
| Residual deviance: 137.01 on 281 degrees of freedom | | | | | |
| AIC: 151.01 | | | | | |
| Type of estimator: AS_mixed (mixed bias-reducing adjusted score equations) | | | | | |
| Number of Fisher Scoring iterations: 8 | | | | | |
| Analysis of Deviance Type II | LR Chisq | Df | Pr(>Chisq) | | |
| Genotype | 41.604 | 1 | 1.118e-10 | **** | |
| Stage_Sex | 5.857 | 5 | 0.3204 | | |

**Table S12.**

Bias-reduced logistic model of CFAV infection status as a function of CFAV-EVE1 genotype and life stage merged with sex variable in Exp12 starting from 3rd instar larvae. Related to Figure 4A.

| Model log10_Infection ~ Genotype * Stage_Sex + (1 \| Line) | | | | | | |
|---|---|---|---|---|---|---|
| Random effects: | | | | | | |
| Groups | Name | Variance | Std.Dev. | | | |
| Line | (Intercept) | 0.0061 | 0.07812 | | | |
| Residual | | 0.08797 | 0.2966 | | | |
| Fixed effects: | Estimate | Std. Error | df | t value | Pr(>\|t\|) | |
| (Intercept) | -2.7053 | 0.0755 | 16.18337 | -35.833 | < 2e-16 | **** |
| Genotype++ | -0.7657 | 0.11025 | 18.23683 | -6.946 | 1.60E-06 | **** |
| Stage_SexAdult_D1_Female | -0.1241 | 0.08562 | 207.0962 | -1.449 | 0.14875 | |
| Stage_SexAdult_D1_Male | 0.52685 | 0.08562 | 207.0962 | 6.153 | 3.87E-09 | **** |
| Stage_SexAdult_D7_Female | -0.4778 | 0.08562 | 207.0962 | -5.581 | 7.44E-08 | **** |
| Stage_SexAdult_D7_Male | 0.49594 | 0.08562 | 207.0962 | 5.792 | 2.55E-08 | **** |
| Genotype++:Stage_SexAdult_D1_Female | 0.06496 | 0.12795 | 207.2264 | 0.508 | 6.12E-01 | |
| Genotype++:Stage_SexAdult_D1_Male | 0.34497 | 0.12625 | 207.328 | 2.732 | 0.00683 | ** |
| Genotype++:Stage_SexAdult_D7_Female | 0.3753 | 0.1271 | 207.3753 | 2.953 | 0.00351 | ** |
| Genotype++:Stage_SexAdult_D7_Male | 0.22228 | 0.12632 | 207.4987 | 1.76 | 0.07995 | . |
| Analysis of Deviance Table (Type III Wald chisquare tests) | | | | | | |
| | Chisq | Df | Pr(>Chisq) | | | |
| (Intercept) | 1284.03 | 1 | < 2e-16 | **** | | |
| Genotype | 48.24 | 1 | 3.77E-12 | **** | | |
| Stage_Sex | 199.654 | 4 | < 2e-16 | **** | | |
| Genotype:Stage_Sex | 13.687 | 4 | 0.008363 | ** | | |

**Table S13.**

Linear mixed-effects model of $log_{10}$-transformed CFAV infection levels as a function of CFAV-EVE1 genotype and life stage merged with sex variable in Exp12 starting from pupae. Related to Figure 4B.

| Model: logit [Infection_status ~ Genotype + Eggs + Generation] | | | | | |
|---|---|---|---|---|---|
| Coefficients: | Estimate | Std. Error | z value | Pr(>\|z\|) | |
| (Intercept) | 5.0874 | 0.7824 | 6.502 | 7.91E-11 | **** |
| Genotype++ | -3.2943 | 0.6725 | -4.899 | 9.63E-07 | **** |
| Eggsaged | -1.4032 | 0.4197 | -3.343 | 0.000828 | *** |
| GenerationF5 | -0.6124 | 0.3715 | -1.648 | 0.099294 | . |
| Null deviance: 246.35 on 235 degrees of freedom | | | | | |
| Residual deviance: 176.65 on 232 degrees of freedom | | | | | |
| AIC: 184.65 | | | | | |
| Type of estimator: AS_mixed (mixed bias-reducing adjusted score equations) | | | | | |
| Number of Fisher Scoring iterations: 4 | | | | | |
| Analysis of Deviance Type III | LR Chisq | Df | Pr(>Chisq) | | |
| Genotype | 53.045 | 1 | 3.26E-13 | **** | |
| Eggs | 13.308 | 1 | 0.000264 | *** | |
| Generation | 2.844 | 1 | 0.091692 | . | |

**Table S14.**

Bias-reduced logistic model of CFAV infection status as a function of CFAV-EVE1 genotype and egg storage condition and generation (females, all collection days merged; normal storage – F4, Exp04 and F5, Exp07; long storage "aged" – F4, Exp17 and F5, Exp18). Related to Figure 5C.

| Model: logit [Infection_status ~ Genotype + Eggs + Generation] | | | | | |
|---|---|---|---|---|---|
| Coefficients: | Estimate | Std. Error | z value | Pr(>\|z\|) | |
| (Intercept) | 5.0874 | 0.7824 | 6.502 | 7.91E-11 | **** |
| Genotype++ | -3.2943 | 0.6725 | -4.899 | 9.63E-07 | **** |
| Eggsaged | -1.4032 | 0.4197 | -3.343 | 0.000828 | *** |
| GenerationF5 | -0.6124 | 0.3715 | -1.648 | 0.099294 | . |
| Null deviance: 246.35 on 235 degrees of freedom | | | | | |
| Residual deviance: 176.65 on 232 degrees of freedom | | | | | |
| AIC: 184.65 | | | | | |
| Type of estimator: AS_mixed (mixed bias-reducing adjusted score equations) | | | | | |
| Number of Fisher Scoring iterations: 4 | | | | | |
| Analysis of Deviance Type III | LR Chisq | Df | Pr(>Chisq) | | |
| Genotype | 53.045 | 1 | 3.26E-13 | **** | |
| Eggs | 13.308 | 1 | 0.000264 | *** | |
| Generation | 2.844 | 1 | 0.091692 | . | |

**Table S15.**

Linear model of $log_{10}$-transformed CFAV infection levels as a function of CFAV-EVE1 genotype and egg storage condition and generation (females, all collection days merged; normal storage – F4, Exp04 and F5, Exp07; long storage "aged" – F4, Exp17 and F5, Exp18). Outliers $<10^{-4}$ were removed from the dataset. Related to Figure S7.
