## Supplementary table 16-17 for "Non-retroviral Endogenous Viral Element Acts as a Stable Antiviral Regulator Across Mosquito Life Stages and Generations"

### Table S16. CFAV-derived EVE sequences tested for detection in this study

| Name | Sequence | Source |
|---|---|---|
| CFAV-EVE1 with flanking regions | AACTACCAGAAACGGTTGTAATGGACAACGAAAAATCATTTTTATCCGGTGAACTTGTTAACTTTTTCAATGAAAACAAAATAACTCCTTATGTAGTAGCTACAGGTCGAAGCGAGATGAACTGTGATATATGTCACAATCAGATTATATTATATACATGTAGTAGTGATAAGCAGAGAATAAAATAAACACAGCGAGCT<br>G GGCTCGCTCTCTTTTATCGTCGACTATCGACCAAACAACACCTATCTTGTTGTACTTTCAATCCGAGAAACTAATAGCAGTAAACCTAAAAAGAACAATCAGTATAGAACATAAAAGTGGCGACGAGGATGGGATCCCAGAAGCACTGGTCCTGAATGGCTTCTTGCACATCTCGCCATGGCACATCCCTGCTCCACG<br>ATCCTATAGCCGCATCATTTCGAACGTTGGCGATGAATTCTTCTTTTGATAAAATACGCGGTCGTAAGTTGCGATTCTTAAAGAACCGGATAAAATGCTTCATGATCGTCCTGTTGACTTTTCCAGAGCTCGAAACTGGTTGAAGGGGGCTACCGCGCCCAAGGCTCTAGCGGCAATGTGCATGAACCTATAAAAAAG<br>GTAGTTCATTTCTAGCAACCCTATGGCAGACACCATGGAGCCAAAATCCATGACATCATGGATCTTTGACCGGGAATGGAGAACTGCGTGACCGGTCTACAACTGAGCGCTCCCTTACCGTGCTAATGGCCCTGTCTCTCGCTACCGTTGTGTTTCTGATCTTCAGTTTGAACGCTATGTACAAGTACGCATGTCAAT<br>ATCACAATCCGAGGAACGGCTACAGAGACTCTGGCGCGAATTTATGGTTTTGGACTGTATCCTTGGCGAGCGCTGGGGGAATCTGGGCGGCGGAGAAAGCTCACCAACCAACGGTGGCAGCTGTCCTAGCCTTTACAATGATAGTGCTCTTTCTTTACATGGAACAAACGAGCGTTAGTATGGAGCTTGAGTTCAT<br>CTCAGCTGGAGAGACTCCCGAAGGAGTGTCAACCGAATATGATGACGGGATAAGTATTCCTGACTTAAAAGTGGCGACGAGTTAAAACAGTGTTTACGCGTGCAAAAGTTCATGTGAAAATCTAAGTGAATTGG | Ref [17] |
| CFAV-EVE2 | GATAAATTTTTACTCTAAGCACCGCATCCAAATCGGGATGTCGGAATGCTTGTTTTAGTTCTTACAGGAGGACATCGTCCAAGCCCTCATTCACATGGCATTGGTGCTTCATGCCCTGTTTGCCTCCATCGACCGGCGAATTCGCTCATTATCTCGACGTGTTACCGCTCTTGAATCTCGACGAACCACAGGAAATCCT<br>ATGACGCTAGCTTTCATCTTGGGCTTTCTCACAGTGTTATGTGGATGCGTGGTGATTGACATGCAAGTTTCAACCGCTAAGGGTACGGAAATTTTTGATGGAAAGACGAACCGGACAGACTACTTGCATCTGTTCAAGTTACCCGCTGATGGTTGCTGGAGTGGCACTCTTGTCACAAAGAAGTGCCCAAAAGTGACT<br>GACTTGGCAAAGGACCTCGAGTCGACAGATTGCGGATCAACGTGGACTGAGTTCACATTGCGGTATCGGAGATGCGTCGTGAAGAAACGTGAAAAGCGATCACGTGAACCCCCGAAAGCGGATCTGCTAGCTGAAATGGAGATCATCGCGTTCAAGACCATTCGCGAAAATAAGACAATTTTCATCGTGGCCCTAT<br>TGTGTGTGGCAATCGCGAAGCGTTGGCCCACCTGGGTCGTGATTCTTCTGGCAATTGGCACTTGGACCACAGTAAAGGGGGAGTTTGTTGAACCGTTATACACGCTGAAAGCGGAACAGATGACCATGCTACAGACCATCGTGAGACCGGAGGAAGGGTACGTCGTGGCCACGCCAAACGGTCTTCTCGAGTTTA<br>AAACAGGACCGGCGGAGATCTATGGGGGGCAATGGCTGCGCGAGTTACTCGCAGATTGCCACGTCAACGCATCCTATTCGACTGACGTGTGTCCCGGTGGATCGCAGCTCAACATGGCTGACATCATGGCCAAGGAACGAGTATGCTCGACCCAACCATACAACCGCGGTTGGGGCACTGGGTGCTTCAAGTGG<br>GGGATTGGATTCGTTGGAACGTGCGTCGAACTCCATTGCGACAGAGGATTCAACGTTTCTTCTATCGCTAGGTCAGCCATTGTCATGAACGTTACGGCTAGCTTCCACTCCGTTAGCGACACACAACAGATGGTTGGCGACATACCGCTCACATTTAGGTTTGCGAAACTCGGAAACGCGGCGATGACCTGCAGGTT<br>GGAGTCTGAGCAGTTGCTGCTTGATTACTATCACGTTACGGGAAGCTCTCACGAGGGGCTGTTTTTACGGTCTCAAGTTGATAGTTGGCCTGGAGTGCACTCTACCGTTAGTGGTCGGCACGGCATGGAGAAAGTTGTTGTGTGGGGAGATGCTAGGTCCAATGAGATTCTAGTAAAAAACGTAATTGAACCTTCGT<br>TGTCCTGGGAAGACGCGATCGCCACCCATGATGGATTCCGTGACATCTCTTTTGTTTGTCAGATTATGCTCGACAAACTAGTCAGTGGTGCTTTTCGAGACTGTCCAGGCCCAAAAACATCCACATTTACCCAAGACGGCTTCGGGTATAGTGGAGTGGTAATAACTACCCTCACCGCTTCTTCTAACGAGACCTGTT<br>CGCTTAGCTTAACGTGCCACGGTTGTCTGCTGCAATCCACAAAAATGATCTTTCTCTCAGGGAAGACCACTTCTCGGGCATTTGTCAAGTGCGGGAACCATACAAGTACCTTGCTCGTTGGGAGCACTTCAGTTTCTATTGAGTGTGCCCTTAACCCTATCTCGCAAGGATGGCGCCTGGCTCGGCACGTGGTGGAT<br>CGGTATCGACGGTTTGGCGTGTCGGGTGTCGCAGGAGTGTGGCAGGATTTGGTTGGAAAGTTTTCGGTTGGTGCCTTCTTCTCCAACACAACCCTTCTGGTCATTTTCGTCCTCGCTGCCCTGATTGACAAACGCATAGCCTTTCTATTTGTTTTAGGGGGGTATTTTTACTATGTGCGGGCTGATCTGGGGTGTGG<br>AATTGACACCACGAGAAAAACCATCTCGTGCGGCAGCGGCGCTTTTGTGTGGAAACACTTGGGCGTTGGCATTTCCAATGACCATGCTGTGGAACTTGAAGATTACTCGTTCACTGACTTGTACATAAAGGATATGTTCAGCTGGACCACGAAACCCTGCTTGATTTGTGAGGACGCTTTGCAGTGCGTAGCCCTGC<br>GCAGAGCTGCTTTCTCTGCTGTTGGGTCGATGGGTCCTGAACAGGTGTATGTAAATGACACATTGGCACGTACATTCAAGTTCTCCGAAACATCAAAGAGAACCATTTCCGTCACCATTAACCTAATTCAGTACAAGTTCTCATCTTATGTTGCACATGGACGCGCTGAGGGTGATTTGGGGCATCTACCAACAATGTT<br>TGGAAGCTATCCCGAAAAAGAGGCTGATAAGGTCATCCGGATTGTCGCATCGCGGCCCGATATTCGCCGACTTTGTGGAAAAGCTGTCTCCTTCCAATTCAAGTTTACGGGCTTCCGGCGAGGCCTGTATGGTTCAAACGTTCAGGTTGAGGTCTCAAAGAACTCATCGACCGAGTGTCCCACGTACCTGGCGGGC<br>GTTGTCGTGAAGAATGGTCGAACTGTCATCACTGACGGCATGTTTTGGATGGAAAGCACAGTGTTGGATGGAGTTGCTCAGATCACATCACTCGAGATGCGTCAATCACACCGCTGTGTGTGGCCCAGGGAATACACCCCGGACACGCTCAGTGACCCTTCAGACCAAGCTTTATTTATTCCACCTGCTTGGGGGG<br>GACCAATTTCGCGAGTGAACCACATAATCGGGTACAAGACACAAACTGACTTTCCGTGGAACGTTAGTGACATCACTCTCATAGAGGGCCCAGCTCCTGGCACAAAGGTCAAGGTGGACTCTAGATGTCATGGGCGAATGCACGCACAGGTGATTGGACCGAATGACACGGAATCTTGGTGTTGCCAGTCATGCAC<br>AAGAATAGTCCACTTTCGAGTCGGTGACCTCCTATATTATCCCATGGAGATCCAACTTGGAACCATGTCTGAAGCGTCAGAGTCGAATTCTAAAATATTTGAGGAACCGATCGGCGAGGAACCAGAACCAACAGTTGACGACATATTGAAGAGATACGGGAAGGCGAACGCGCAATCGGATTTCCGGAGGGTGAGCT<br>CACGTGCAGGAGTTGGGTTCGATCGCTCCCTCCTAAATCTCCTGTGCCTCGCTATCTCATTGCAGCTTATCGGGGCAAAAACCAGAACATCGACACTAACTCGTCTATTTCTGACGATTCTAGCAATGGCGCTCTTTGGACTCCCCAATTTGTTTTCATCTGTGGGGCTATCTGCTTGGGTGTTGCTAGTGGCGTCCT<br>CATCATCGCAACCGCAAGATCTATTGATGAATCTATGGATCATACTACAGACTGGGAGCTCAGCAGTCCTGCTGTTGGGGTACATGATCAGGAAGAAACTCTCAATAGTGCTAGGGGTCCACCACCTGGTGACTCTGATGTGCGTACAGT | Ref [17] |
| CFAV-EVE3 with flanking regions | TGTACGACCAGCACCCCGTCGCCCAGCTGTTAAGATGTATGTTGCTGTGTTCATCTGCCTTTGCACTAAGGCGGTCCATCTGGAGCTTGTATCCGACCTCACAACCGAACGATTTATACAAGCTCTGCGTCGATTTGCATCCAGGAGGGGCAAGCCCAAGGACATGTACTCCGACAACGGTACTAACTTCGTAGGG<br>GCATGCAACAAGCTGCGAGACTTGAACGATTTGTTAAAAAGTCCTGCGCATAAAGAGAAGATAGCTTATCATTGCTTGGAAGAAGGGACCCAATGGCACTTTAATCCGCCAAGTGCCCCACACTTTGGAGGTTTATGGGAGGCGGCAGTGCGGTCAGCTAAGGGGAGAGGATGGGAAGAGTTTTCGGCTTCAAACG<br>ATCAGGATACTTGAAGGACATTGCTAGCAGGTGGTAAACCAACCGATGTCACGGAGAACTTGAACTGTGATTCGACCAGGCGGTTCGTGACATGGCACCCAGGAAAAGGAAAGACGCGCAAGGTTATTGTCGAAGAAACCAAGAAGAACTATGATTCCAACCAACGGACCGTGATCTTAACACCCACCCGGGTAGT<br>CATGGCTGAGGTCGTTGAGGCCTTGAAGAACTCAGGCATGAGGAGTGATAAGAACCTGTCGTACTGCACCAGAAACCTAATAACGGTTGCATGTCACGCAACGTTCACGAAATTTGTTCTTAGCCATGGCGCCAAGAAAGTGCGGGTTGCCATGATCATCATGGATGAGTGCCATTTCATGGATCCTATGTCCATTG<br>CTGCGCGCGGGGTCATGGAACATCTCCACAGCCAAGGAACGAAATTGATTTACCTTAGTGCAACACCACCCGGCCACGCGCCTGATACAGGCTCGAATTATGCAATAAGTGACCAATCCATTAGCTTCCCAACTTGGTTGAGTCCAGCGTGGATTGGAAACGTTCAAAAATCCGTGGGGGCGAAGAAGACCATTTT<br>ATTCGTCCCATCGCACAACCAAGCTAACACCCTGGCTAGTGCTATTCCTGGATCCGTTCCATTGCATCGAGCAAACTTTTCCACTAACTATGCACAAGCAGGTGATGCAGCCACCGCCCTGGTAATCTCCACCGACATCTCAGAAATGGGAGCGAATTTGGGCGTCGATCTGGTTATAGACACTCGAAGAGCTTTGC<br>GACCCCTTGTGGATTCTGCCACGCGTGTGAAGCTGGTTGAAACGAACATAACCACATCATCCATGATCCAGAGGAGAGGGAGGACGGGTAGACGGGAACCAGGAACGTATGTGTATCCTATTGATAGCCAAACGGAGGAAAATCCAGTGTCATGGGTGTGCTGGCCTGAAGCCCAAATGATCCTGGACCAGCTCG<br>GCATGACTTTTATGCTAGAGGAAGCCGCATACAGCCAACCTCCGGGGAGATTTACCCTAGTGGGAGAGGATAGAATGCGATTTTCAAAGCTTATGGACCGGGATGACATTCCCATTTGGTTGGCGTGGCACTGGGCAGAAGCTGGAGACCGACGGCACTCTGCCCTGTTTCAGGGCGAAGGAACCGGAAAGATCA<br>TCGAAAACCGATTCGGCAAACAAGAGTATCGACCACAATACGTTGATGACCGCTTTGAGAGCATTGAGTGGGAGACGCGAAAGGTATCCATTGATTTCTACATGAACTGCCGCGGGGGGCCTAGCCTCTATGAGTTTTTCACAGTGATTGATTGGACCGATATATGGAGGAGAACGACATCTGCTCTCTGGGACTTG<br>AGTGACGTCATGAATGGAGAACTGCGTGACCGGTCYACAACTGAGCGCTCCCTTACCGTGCTAATGGCCTTTGTCTTGGGCGTTTCAATTGTGCTCTCCTGTTTCATCGCTGTCTGGGCATTGTGTTTCCTATTCTCCTTGTTTAGGCCGCGAAAGGTAACTTATGAACAAATGCCATCTAGTGACCCACTTAGCGGT<br>GGTGTCATGGTGTCAACCCCTAGCCTCCTTTATTATATGGGAGTCCCCTTGGGGTTCTGCGTCGTGAAGAAACGTGAAAAGCGATCACGTGAACCCCCGAAAGCGGATCTGCTAGCTGAAATGGAGATCATCGCGTTCAAGACCATTCGCGAAAATAAGACAATTTTCATCGTGGCCCTACTGTGTGTGGCAATCGC<br>AAAGCGTTGGCCCACCTGGGTTGTGATTCTTCTGGCAATTGGCACTTGGACCACAGTAAAGGGGGGAGTTTGTTGAACCGTTATACACGCTGAAAGCGGAACAGATGACCATGCTACAGACCATCGTGAGACCGGAGGAAGGGTACGTCGTGGCCACGCCAAACGGTCTTCTCGAGTTTAAAACAGGACCGGCGGA<br>GATCTATGGGGGGCAATGGCTGCGCGCTTCTTCTAACGAGACCTGTTCGCTTAGCTTAACGTGCCACGGTTGTCTGCTGCAATCCACAAAAATGATCTTTCTCTCAGGGAAGACCACTTCTCGGGCATTTGTCAAGTGCGGGAACCATACAAGCACCTTGCTCGTTGGGAGCACTTCAGTTTCTATTGAGTGTGCCC<br>TTAACCCTATCTCGCAAGGATGGCGCTTGGCTCGGCACGTGGTGGATCGGTATCGACGGTTTGGCGTGTCTGGTGTTGCAGGAGTGTGGCAGGATTTGGTTGGAAAGTTTTCGGTTGGTGCCTTCTTCTCCAACACAACCCTTCTGGTCATTTTCGTCCTCGCTGCCTTGATTGACAAACGCATAGCCTTTCTATTG<br>GTTTTAGGGGGGTACTTTTACTATGTGCGGGCTGATCTGGGGTGTGGAATTGACACCACGAGAAAAACCATCTCGTGCGGCAGCGGCGCTTTTGTGTGGAAACACTTGGGCGTTGGCATTTCCAATGATCATGCTGTGGAACTTGAAGATTACTCGTTCACTGACTTGTACATAAAGGATATGTTTAGCTGGACCAC<br>GAAACCCTGCTTGATTTGTGAGGATGCTTTGCAGTGCGTAGCCCTGCGCAGAGCTGCTTTCTCTGCTGTTGGGTCGATGGGACCCGAACAGGTGTATGTAAATGACACATTGGCACGTACATTCAAGTTCTCCGAAACATCAAAGAGAACCATTTCCGTCACCATTAACCTAATTCAGTACAAGTTCTCATCTTATGTT<br>GCACATGGACGCGCTGAGGGTGATTTGGGGCATCTACCAACAATGTTTGGAAGCTATCCCGAAAAAGAGGCTGATAAGGTTATCCGGATTGTCGCATCGCGGCCCGATATTCGCCGACTTTGCGGGAAAGCTGTCTCCTTCCAATTCAAGTTTACGGGCTTCCGGCGAAGCCTGTATGGTTCAAACGTTCAGGTTG<br>AGGTCTCAAAGAACTCATCGACCGAGTGTCCCACGTACCTGGCGGGCGTTGTCGTGAAGAATGGCCGAACTGTCATCACTGACGGCATGTTTTGGATGGAAAGCACAGTGTTGGATGGAGTTGCTCAGATCACATCACTCGAGATGCGTCAATCACACCGCTGTGTGTGGCCCAGGGAATACACCCCAGACACGC<br>TCAGTGACCCTTCAGACCAAGCTTTATTTATTCCACCTGCTTGGGGGGGACCAATTTCGCGAGTGAACCACATAATCGGGTACAAGACACAAACTGACTTTCCGTGGAATGTTAGTGACATCACTCTCATAGAGGGCCCAGCTCCTGGCACAAAGGTCAAGGTGGACTCTAGATGTCATGGGCGAATGCACGCACAG<br>GTAATTGGACCGAATGACACGGAAACTTGGTGTTGCCAGTCATGCACAAGAATAGTCCACTTTCGAGTTGGTGACCTCCTATATTATCCCATGGAGATCCAACTTGGAACCATGTCTGAAGCGTCAGAGCCGAATTCTAAAATATTTGAGGAGCCGATCGGCGAGGAACCAGAACCAACAGTTGACGACATATTGAA<br>GAGATACGGGAAAGCGAACGCGCAATCGGATTTCCGGAGGGTGAGCTCACGTGCAGGAGTTGGGTTCGATCGCTCCCTCCTAAATCTCCTGTGCCTCGCTATCTCATTGCAGCTTATCGGGGCAAAAACCAGAACATCGACACTAACTCGTCTATTTCTGACGATTCTAGCAATGGCGCTCTTTGGACTCCCCAATT<br>TGTTTTCATCTGTGGGGCTATCTGCTTGGGTGTTGCTAGTGGCGTCCTCATCATCGCAACCGCAAGATCTATTGATGAATCTCTGGATCATACTACAGACTGGGAGCTCAGCAGTCCTGCTGTTGGGGTACATGATCAGGAAGAAACTCTCAATAGTGCTAGGGGTCCACCACCTGGTGACTCTGATGTGCGTACAG<br>TTCCTGTTCTCCGCCGTCGATCGCTATCAGAAATATCTCTACGGTCTCCTAGAACTCATGGCTTCGGTGGTCCTGCTGAGCGCCTACAAGAGCGTCCTGCAGGCTCTCCCTCCAGAGGTTCTATGTTTCAGCCTGATAATGGGGTGGAAGAGCGGCCCTGTCTCTCGCTACCGTTGTGTTTCTGATCTTCAGTTTGAA<br>CGCTATGTACAAGTACGCATGTCAATATCACAATCCGAGGAACGGCTACAGAGACTCTGGCGCGAATTTATGGTTTTGGACTGTATCCTTGGCGAGCGCTGGGGGAATCTGGGCGGCGGAGAAAGCTCACCAACCAACGGTGGCAGCTGTCCTAGCCTTYACAATGATAGTGCTCTTTCTTTACATGGAACAAACG<br>ARYGTYAGTATGGAGCTTGAGTTCATCTCAGCTGGAGAGACTCCCGAAGGAGTGTCAACCGAAAATGATGACGGGATAAATATTCCTGACTTAAAAGGGCGCTATGGGGAGGATGGGATCGTCGTGGGGGCGGCCTCATCGTCTGGCTACCTACCTGAGCTCGTATTTGTGTTCCTCCTAGGCTTTGCCGTGACGT<br>CCACCTCATACTTTTTGGGGGCACTCTACCTGCTCATCGCAACCAGCACTAACCTTCCCGTGACCATAATACGGATGCTGCGTGTGAAGCTCGCTGCTAACAACCGATCTGATGATCTCTTGGGACTCGGCGGACCTATTGAAACTGATCTCCAGACCAGTTTCAAGACATTCCCAATGGAGTGTACCGCATCGTT<br>GTTCGCAGCTTATTTGGCGACCGGCAACGAGGGGCTGGCTTTTCGAAAAATGGGGTATTTCACACTCTCATGCACGTCACTCGAGGTGAACCAGTTAAATGGCGTGGGCGAGTGTGTCGTTCTATTGCCACGGAAGAATAGTGCCCAAAACCCAAAACTACCCCGGTAAGGATCTGGACGCTCAACCACTGGAAGG<br>TCTCAATCGGATCCCCTTATTCCACTCACAATCAATCCGTTCGAGTGTTGAGGAGCCCAAACGAAATCGAAGGAGGTAAATTCCCCGGTTCCCCAACATGGTTCATGGCAGAGAGAGGTTAATGTGAAAAAAGGTAAAAAAAGTGAAATAAAGAAACTTTGGACGCTTACATCTCAAAAACAGTTTTACATGAAACTAT<br>ACTTTTTCGGTAAGGATTTGCAACAGTGGAAAATCTATTTCTGAGCTTTGCAATTTTCGAGAACTTTCTCCGAGTTGGTAAACATTGCCACATTATCGAACATCATCAATAAAACAAACTTGTTTTTTTTTGTTGAAAATAACTGAT | Ref [21] |
| CFAV-EVE4 with flanking regions | ACCTGCGTTGACCCTAGGCGGCTAATTCAGGAGTGAGTCCGTGAAGCCCTGTACCCCAGAACGACCACCAAGTGCGCCTGGAAACCCAGGCTTTCCAGTGAATCCACGGATCACTGAAGCCTCGAGTCCGTCACCACACTCGAGGGGCGCACGGGACAGGCACGAGGTCGACGAGAGGCTGCAGACGAAGGAG<br>GAGAAACGAGAAGGAAGGAGTCAACCGCCCCTGGCCCGGATAGCGGAAGTAGTTCTAGTGGTCAGTGAAGGAGCAGCGGAGAAGGTGCTGCAGTGCGCGGGGCCCCGAAGAAGTGAAGGAATTGTAAGGTAGGATAGGATAAGAAAGTGGGAGGCGTCGTCCGGTGTTGAGGACGGACGCAAGTAAAGAGAA<br>GGAAAGATTAGGAGAGAGYGGGGAGTAGAAGAAGTAGGACGTCAAGGAAGTGGAAATAAAAGTGTACCCGAGACCACGAAATAGAGGGAGTTTTACTAGAGTCGATCTGTAAGTGTTTTCTTTCGCGGAGTGTCACGAGCGTAGCGCCGACCCTGAGGCTTGGAGACGGGGGTCTCATCTAGGCTGGGAGTAACA<br>CGGCCTGCAAGTTACACTTTGGCGCCCAACGTGGGGCTGGCTTTTCGAAAAATGGGGTATTTCACACTCTCATGCACGTCACTCGAGGTGAACCAGTTAAATGGCGTGGGCGAGTTGTCGTTCCTCACTCTGGTAGTGCCTTCGCGCGACGTGGTGTCGTATGGTGGACCTTGGCAGTTGGACACCCCCACTACCA<br>CGGAAGACCTCGTGCTGATGGCTTGCAAACCGGACAAGACCATTGAGTACCATCGGTACAGGCCGGGAGTCATGAGTATTGACGGTGAACCTGTCATGTTCATTAGTGATGATTTCGGAAAAGGATCTTCTGGCTCACCGTTCTTCATCAATGGCGAGCCCGTCGGGTTCTATGGTTTTGGGTTCTATGTGAACGGT<br>GTTTATCGCTCAACAGTAGCAGGTGGTAAACCAACCGATGTCACGGAGAACTTGAACTGTGATTCGACCAGGCGGTTCGTGACATGGCATCCAGGAAAAGGAAAGACGCGCAAGGTTATTGTCGAAGAAACCAAGAAGAACTATGATTCCAACCAACGGACCGTGATCTTAACACCCACCCGGGTAGTCATGGCTG<br>AGGTCGTTGAGGCCTTGAAGAACTCAGGCATGAGGAGTGATAAGAACCTGTCGTACTGCACCAGAAACCTAATAACGGTTGCATGTCACGCAACGTTCACGAAATTTGTTCTTAGCCATGGCGCCAAGAAAGTGCGGGAACCATACAAGTACCTTGCTCGTTGGGAGCACTTCAGTTTCTATTGAGTGTGCCCTTAA<br>CCCTATCTCGCAAGGATGGCGTCTGGCTCGGCACGTGGTGGATCGGTATCGACGGTTTGGCGTGTCTGGTGTTGCAGGAGTGTGGCAGGATTTGGTTGGAAAGTTTTCGGTTGGTGCCTTCTTCTCCAACACAACCCTTCTGGTCATTTTTGTCCTCGCTGCCCTGATTGACAAACGCATAGCCTTTCTATTGGTTT<br>TAGGGGGGTATTTTTACTATGTGCGGGCTGATCTGGGGTGTGGAATTGACACCACGAGAAAAACCATCTCGTGCGGCAGCGGCGCTTTTGTGTGGAAACACTTGGGCGTTGGCATTTCCAATGATCATGCTGTGGAACTTGAAGATTACTCGTTCACTGACTTGTACATAAAGGATATGTTTAGCTGGACCACGAAA<br>CCCTGCTTGATTTGTGAGGATGCTTTGCAGTGCGTAGCCCTGCGCAGAGCTGCTTTCTCTGCTGTTGGGTCGATGGGTCCTGAACAGGTGTATGTAAATGACACTTTGGCACCCAACGTGAATTGTCTTTGGCATTCCGTGCTAGGGAAACACTTTAGATTCTGATAACTTTTCTTTGCGGAGGTCTCTGGGAAAACA<br>TCCCGAAGTCACTTACGGAATTGACGTCACGAACTACTTTGGAGTTGTTCAGAACTTGTAATCATTGTTACCATAACGGAGGTCTTGGAAGTCCCAAGTGGCAAGATAGCTACCAATCATTGTTGAGAACGAATTTAAAACAGAATCGGAGTAATTTGTTAAATTTTGCGATAGAATTGCGGAAGTTATCTTGATCTGTG<br>TGTTTCGGAGAAGAGTGTTGAGAAACACCGAGCAAATCTAGGAACTATATTTCAATTTTAACCGAGTGAACCAATCAGACTTGGTCAGCATTTTTCTAGGATTTTCAAATTATTTAGAGCTTTTGCTAAGAACCGTGAACAAATTTTAGGTACCTTTTCAAATTAATTTAGCTTTCGAAAAACGAAAGCGTTTATAAAACTA<br>GGTATTTTTTTCAAATTTTACAGCTTTGTGCTGAAATAGGTTT | Ref [21] |
| CFAV-EVE5 with flanking regions | AAAACATCATTTGGAATTTACAAAACATAAAAATAAATTAGTATCAAATAAAAGAGCATGTGGCTCGACAACGACAGTTCGTTCAGTTCGTTCAGATAGTTCAGTCTTAAGATAGTTTTTTTATAATTAAGTTTTAGTTCCAAATTTATTAACTGGCCCCCGAATAGGGACCAGCGCGAGTGTAGTTTAATTGTGTTGCGAG<br>TGTTAAAACATCGTGAGCATTATCCGAAAAATTCAACTTTACCGCAGCACAAGTGGCGACCGCTACTAACGCAGCAGACGTGGAGAAAGCGCCCCTAGTAACCACCCTATAGGAAAAACGGGATCATCGCATCTTCGCTGTGTAAGCAAGCCTGAATAAAGCACAAAGTGACTTAATCACTGTGCGACATAATTGACA<br>AATCCATAACACATCATTTTAAATAATTCTCACGAAATAGTTTAGCAACATGCAGTCACTTTGCACTCTTCAAGTGCCAACCATCATTGCACCCTGTTTGGGTGTTGCTAGTGGCGTCCTCATCATCGCAACCGCAAGATCTATTGATGAATCTATGGATCATACTACAGACCGGGAGCTCAGCAGTCCTGCTGTTGGG<br>GTACATGATCAGGAAGAAACTCTCAATAGTGCTAGGGGTCCACCACCTGGTGACTCTGATGTGCGTACAGTTCCTGTTTTCCGCCGTCGATCGCTATCAGAAGTATCTCTACGGTCTCCTAGAACTCATGGCTTCGGTGGTCCTGCTGAGCGCCTACAAGAGCGTCCTGCAGGCTCTCCCCCCAGAGGTTCTATGTT<br>TCAGCCTGATAATGGGGTGGAAGACGGCAAGCAAAAGATAGGATTACAGACCTCCTTGCTTGCTATGAGAACCTAATTGCTCGTTACACATGTTGCATATATTATTGGTATTGGTTCGATCGGAGCTACCACCTGTGGTGGATACGGACCATATTCAATGCACGGTCAGCATAACCGCAATCAAGTCCAAAGTTAAAC<br>AAACTCGGACCATTGCTGTTTATGCTGACTGCAGTGAAAAGTGTCACGTAGGCACACCAACCGCGAGTAGCGTTAAGCATAGAGAAAAATGTTAAAATTCATTGTTGTGTCGACAATCAACTGAGTTGGGTCAATTTATTACAGTGACAGTCCAGATGTTACATGGCGATCCTGCCATTTCTCCTGGGTGACTGAAAAA<br>AGATAACAAATAGTTAATAACCAGTGAAAAACATGGTTCTGCGAACAACACTATTAGTGAATATAAAGGAAAAAAAGTATAACAGAGCGTA | Ref [21] |

**Table S17. Primer sequences used in this study**

| Target name | Purpose | Forward primer sequence | Reverse primer sequence |
|---|---|---|---|
| CFAV-EVE1 | CFAV-EVE1 detection and genotyping | CTTATGTAGTAGCTACAGGTCGAAGCGAG | GCAATGGCCAAATGCTGCTGCGAG |
| CFAV-EVE2 | Detection of CFAV-EVE2 | CCTCATTCACATGGCATTGGTG | AGGAACTGTACGCACATCAGAG |
| CFAV-EVE3 | Detection of CFAV-EVE3 | AGCCCAAGGACATGTACTCC | AAAATGGTCTTCTTCGCCCC |
| | | ACTCTCAATAGTGCTAGGGGTC | AGTTTTGGGTTTTGGGCACT |
| CFAV-EVE4 | Detection of CFAV-EVE4 | CCCGGATAGCGGAAGTAGTT | GACAATAACCTTGCGCGTCT |
| | | AAGTACCTTGCTCGTTGGGA | GATGTTTTCCCAGAGACCTC |
| CFAV-EVE5 | Detection of CFAV-EVE5 | ACTTTACCGCAGCACAAGTG | GCTCCCGGTCTGTAGTATGA |
| | | ACGGTCTCCTAGAACTCATGG | GGTAGCTCCGATCGAACCAA |
| *Ae. aegypti* s7 | Detection of s7 | GGGACAAATCGGCCAGGCTATC | TCGTGGACGCTTCTGCTTGTTG |
| CFAV NS3 | qPCR for CFAV | ACACGAGTGAAGCTGGTTGA | ACATACGTTCCTGGTTCCCG |
| *Ae. aegypti* rp49 | qPCR for rp49 | ACAAGCTTGCCCCCAACT | CCGTAACCGATGTTTGGC |
